# Targeting Synaptic Vesicle Endocytosis in Nociceptors Provides Sustained Pain Relief

**DOI:** 10.64898/2026.06.17.732993

**Authors:** Raquel Tonello, Maria Fernanda Pessano Fialho, Thomas Payne, Elisa Damo, Martina Chieca, Romina Nassini, Francesco De Logu, Wendy L. Imlach, Colin W. Pouton, Nigel W. Bunnett

## Abstract

Endocytosis replenishes synaptic vesicle (SV) pools that are required for persistent transmission of chronic pain signals within nociceptive spinal circuits. The nociceptor-specific contribution of SV endocytosis to pain and the therapeutic potential of endocytosis inhibitors are unclear. We identified SV endocytosis in nociceptors as a critical driver of ongoing pain and developed a gene-based strategy to target this mechanism. Nociceptor-specific adeno-associated virus-mediated knockdown of adaptor-associated kinase 1 (AAK1) or dynamin 1 (Dnm1) in dorsal root ganglia Nav1.8-positive neurons inhibited postoperative and neuropathic hypersensitivity without affecting baseline mechanical or thermal sensitivity, locomotion or spontaneous behavior. Electrophysiological recordings from spinal neurons combined with optogenetic activation of nociceptor afferents showed that AAK1 or Dnm1 downregulation blocked the sustained synaptic transmission between nociceptors and dorsal horn neurons by disrupting SV recycling and reducing neurotransmitter release probability. Lipid nanoparticle (LNP)-encapsulated CRISPR/dCas9-repressor mRNA constructs (dCas9-R) were engineered to achieve sustained and reversible transcriptional and epigenetic repression of *Aak1* or *Dnm1* following intrathecal delivery. LNP-mediated gene modulation produced sustained downregulation of *Aak1* or *Dnm1* mRNA in sensory neurons and resulted in robust and long-lasting analgesia in preclinical models of postoperative, inflammatory, neuropathic and osteoarthritis pain without impairing acute nociception or locomotor activity. Mechanistically, targeting endocytic machinery disrupted SV recycling at nociceptor terminals, thereby reducing excitatory neurotransmission within spinal pain circuits. Together, these findings establish presynaptic endocytic regulation as a convergent mechanism underlying chronic pain and demonstrate the translational potential of LNP-delivered CRISPR/dCas9-R as a durable, non-opioid pain therapy that surmounts inherent redundancy of pain signaling mechanisms.

**One Sentence Summary:** Synaptic vesicle endocytosis in nociceptors is a critical mechanism driving ongoing pain and targeting this process with intrathecal LNP-delivered CRISPR/dCas9-mediated gene repression produces durable, non-opioid analgesia across multiple chronic pain models.

## Introduction

Chronic pain is a debilitating condition that afflicts >20% of individuals worldwide, imposing a substantial public health burden(*1, 2*). Despite intensive effort, safe and effective treatments for chronic pain remain elusive. Common pharmacological therapies, including nonsteroidal anti-inflammatory drugs, glucocorticoids and paracetamol, offer only partial relief and can have serious deleterious effects on the gastrointestinal, cardiovascular, hepatic and renal systems(*3–5*). Opioids have life threatening side effects that are exacerbated by addiction and dose escalation to counter tolerance(*6, 7*). These challenges underscore the pressing need for safe and effective therapies for chronic pain.

Painful stimuli in the periphery are transmitted to the brain by a common pathway. Painful somatic or pathological stimuli are detected by the peripheral terminals of primary afferent neurons (nociceptors) with cell bodies within the dorsal root ganglia (DRG). Nociceptive signals trigger the release of neurotransmitters, including glutamate, substance P (SP) and calcitonin-gene-related peptide (CGRP), from the central terminals of these neurons in the dorsal horn of the spinal cord, where the input is encoded and transmitted to the brain for processing of the sensation of pain(*6, 8, 9*). We have previously reported that the endocytic retrieval of synaptic vesicles (SVs) in the central terminals of primary afferent neurons maintains the releasable pool of SVs and is required for ongoing synaptic transmission to spinal neurons. Dynamin (Dnm) 1, adaptor-associated protein kinase 1 (AAK1), endophilin A1 and synaptojanin were found to be required for the synaptic transmission of painful signals(*10, 11*). The local (intrathecal) administration of small molecule inhibitors or siRNA/shRNA targeting these endocytic mediators depleted the releasable SV pool, blocked synaptic transmission in nociceptive spinal circuits, and blunted pain-like behavior in multiple preclinical pain models, without affecting normal behavior(*10, 11*). However, the specific role of endocytic mediators within nociceptors, the peripheral sensory neurons that initiate pain signaling, remains poorly defined. Given the redundancy of targeting pain receptors and ion channels, disrupting a common pathway of pain transmission may offer broader therapeutic efficacy, positioning endocytic mediators as promising targets for translational pain therapy.

Gene therapy has emerged as a promising approach for chronic pain treatment by enabling precise, tissue-specific modulation of pain-related targets(*12, 13*). Advances in clustered regularly interspaced short palindromic repeats (CRISPR) technology have further expanded this approach, with CRISPR-based strategies demonstrating therapeutic efficacy in multiple animal models of human diseases(*14–16*), supporting their potential application in pain management. Lipid nanoparticles (LNPs) provide an efficient, non-viral delivery system for gene editing and are clinically validated through their widespread use in mRNA vaccines(*17–19*). LNPs can effectively deliver CRISPR-based mRNA components to enable either transient transcriptional modifications via dead Cas9 (referred to here as dCas9-R, also known as CRISPRi) or permanent genomic modifications via Cas9 of target genes (*20, 21*). In the context of pain management, LNP-encapsulated CRISPR offers a means to achieve targeted and prolonged modulation of pain-related genes in nociceptive neurons, with the potential for long-lasting analgesia and minimal off-target effects.

Here, we used targeted genetic strategies to selectively disrupt SV endocytosis in nociceptors and demonstrated that the endocytic mediators AAK1 and Dnm1 are required to sustain pain hypersensitivity by maintaining SV recycling and synaptic transmission in nociceptive circuits. Building on this mechanism, we developed a novel therapeutic approach for pain treatment that uses LNPs to locally deliver mRNA CRISPR/dCas9-R constructs with a single guide RNA (sgRNA) to downregulate the genes encoding *Aak1* and *Dnm1*. LNP dCas9-R AAK1 and LNP dCas9-R Dnm1 effectively reduced pain-like behavior in preclinical models of postoperative, inflammatory, neuropathic and osteoarthritis pain, while not affecting normal behaviors. Together, these findings establish SV endocytosis in nociceptors as a key driver of ongoing pain and support LNP-mediated CRISPR/dCas9-R gene modulation as a promising, non-opioid therapeutic strategy for chronic pain.

## Results

### Targeting AAK1 or Dnm1 in nociceptors results in cell-type specific knockdown

We have previously reported that AAK1 and Dnm1 are required for sustained synaptic transmission in nociceptive spinal circuits(*10, 11*). To investigate the role of AAK1 and Dnm1 specifically in nociceptors, we used a Nav1.8^Cre^ driver to express shRNA for the selective silencing of *Aak1* or *Dnm1* in nociceptors using a Cre-dependent adeno-associated viral vector (AAV). We generated an AAV-PHP.S serotype for efficient infection of peripheral neurons with loxP-flanked shRNAs to *Aak1*, *Dnm1* or their respective *scrambled controls (Scr)* and a GFP reporter. The virus was administered by intrathecal injection (i.t., 5 µL, 1×10¹² v/g, L4/L5) into Nav1.8^Cre+^ or control (Nav1.8^Cre-^) mice (**Fig. 1a**). After 3 weeks, *Aak1* and *Dnm1* mRNA were localized in DRG (L4/L5) using RNAScope *in situ* hybridization (**Fig. 1b**). In DRGs from Nav1.8^Cre+^ mice treated with control AAV *Scr* shRNA, *Aak1* or *Dnm1* mRNA were detected in both *Scn10a* positive and negative (Nav1.8+ve; -ve) neurons (**Fig. 1c, h**). Selective knockdown of *Aak1* or *Dnm1* mRNA in Nav1.8+ve neurons in the DRGs was achieved with AAV *Aak1* shRNA or AAV *Dnm1* shRNA, respectively, after 3 weeks, compared with AAV *Scr* shRNA (**Fig. 1dI, e, iI, j**). The expression of floxed GFP reporter gene in Nav1.8+ve neurons (**Fig. 1dII, iII**) and the lack of *Aak1* or *Dnm1* mRNA downregulation in Nav1.8-ve neurons and in spinal cord, support cell-type selective targeting (**Fig. 1f, g, k, l** and **Fig. S1a, b**).

**Figure 1.**
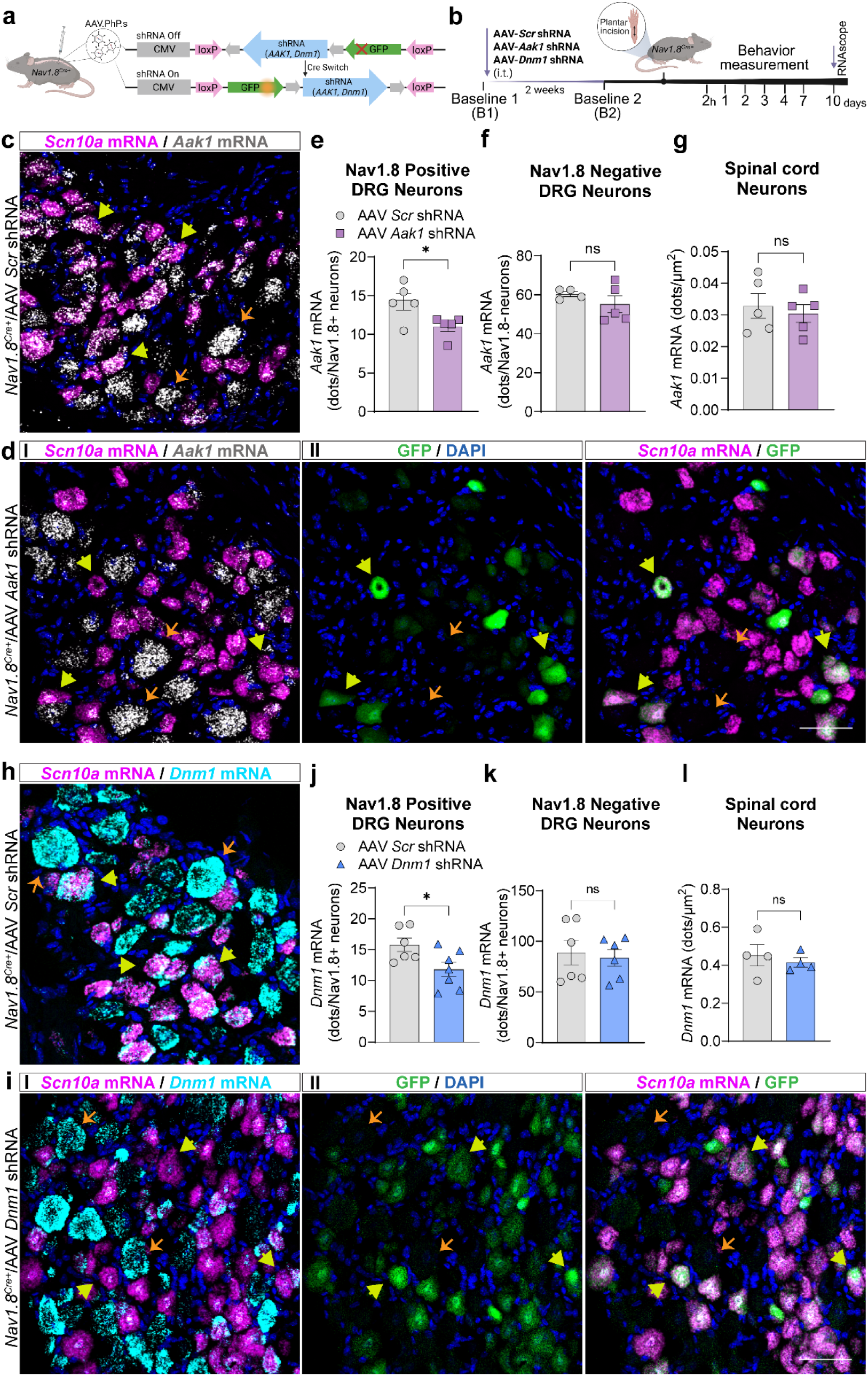
Selective knockdown of AAK1 and Dnm1 in nociceptors. Schematic of AAV loxP-flanked shRNA to *Aak1* or *Dnm1* vector pre- and post-Cre switch (a). RNAScope experimental timeline following the postoperative pain model (b). Localization (c, dI, dII) and quantification (e, f, g) of *Aak1* mRNA expression in Nav1.8+ve and Nav1.8-ve neurons in DRG and spinal cord neurons 3 weeks after intrathecal (i.t.) injection of AAV *Aak1* or *scrambled* control (*Scr)* shRNA in *Nav1.8^Cre+^* mice. n=4-5 mice per group. Localization (h, iI, iII) and quantification (j, k, l) of *Dnm1* mRNA expression in Nav1.8+ve and Nav1.8-ve neurons in DRG and spinal cord neurons 3 weeks after i.t. injection of AAV *Dnm1* or *Scr* shRNA in *Nav1.8^Cre+^*mice. n=6-7 mice per group. GFP-positive cells in Nav1.8+ve neurons (dII, iII). Two-tailed unpaired t-test, *P<0.05. Yellow arrows show expression with Nav1.8+ve neurons, orange arrows show Nav1.8-ve neurons. Scale bar: 50 µm.

### Selective knockdown of AAK1 or Dnm1 in nociceptors prevents postoperative pain

The effects of AAK1 and Dnm1 knockdown in nociceptors were evaluated in a preclinical model of postoperative pain induced by plantar incision into the left hindpaw of Nav1.8^Cre+^ or control (Nav1.8^Cre-^) mice. Two weeks prior to the incision, AAV *Aak1, Dnm1* or *Scr* shRNA was administered by i.t. injection. Withdrawal responses of the incision (left, ipsilateral) and non-incision (right, contralateral) hindpaws to von Frey filament and radiant heat stimulation were assessed from 2 hours to 7 days to evaluate mechanical allodynia and heat hyperalgesia, respectively. (**Fig. 2a**). Plantar incision reduced both the withdrawal threshold to von Frey filaments and the withdrawal latency to heat in the ipsilateral paw for at least 4 days, consistent with mechanical allodynia and heat hyperalgesia, with a resolution on day 7 (**Fig. 2b-e**). AAV *Aak1* shRNA or AAV *Dnm1* shRNA completely prevented mechanical allodynia for at least 4 days and substantially reduced heat hyperalgesia for 3 days post-incision, compared to AAV *Scr* shRNA in both male and female Nav1.8^Cre+^ mice, with no effects in Nav1.8^Cre-^ mice. No differences were observed between male and female mice (**Fig. S2**; graphs in **Fig. 2** show combined data). Importantly, AAK1 or Dnm1 knockdown did not alter baseline mechanical or heat sensitivity before the incision or affect the withdrawal responses of the contralateral (non-incision) paw to mechanical and heat stimuli, demonstrating that the downregulation of AAK1 or Dnm1 in nociceptors does not affect normal behavior (see **Fig. 2** baseline B2, and **Fig. S3a-d**).

**Fig 2.**
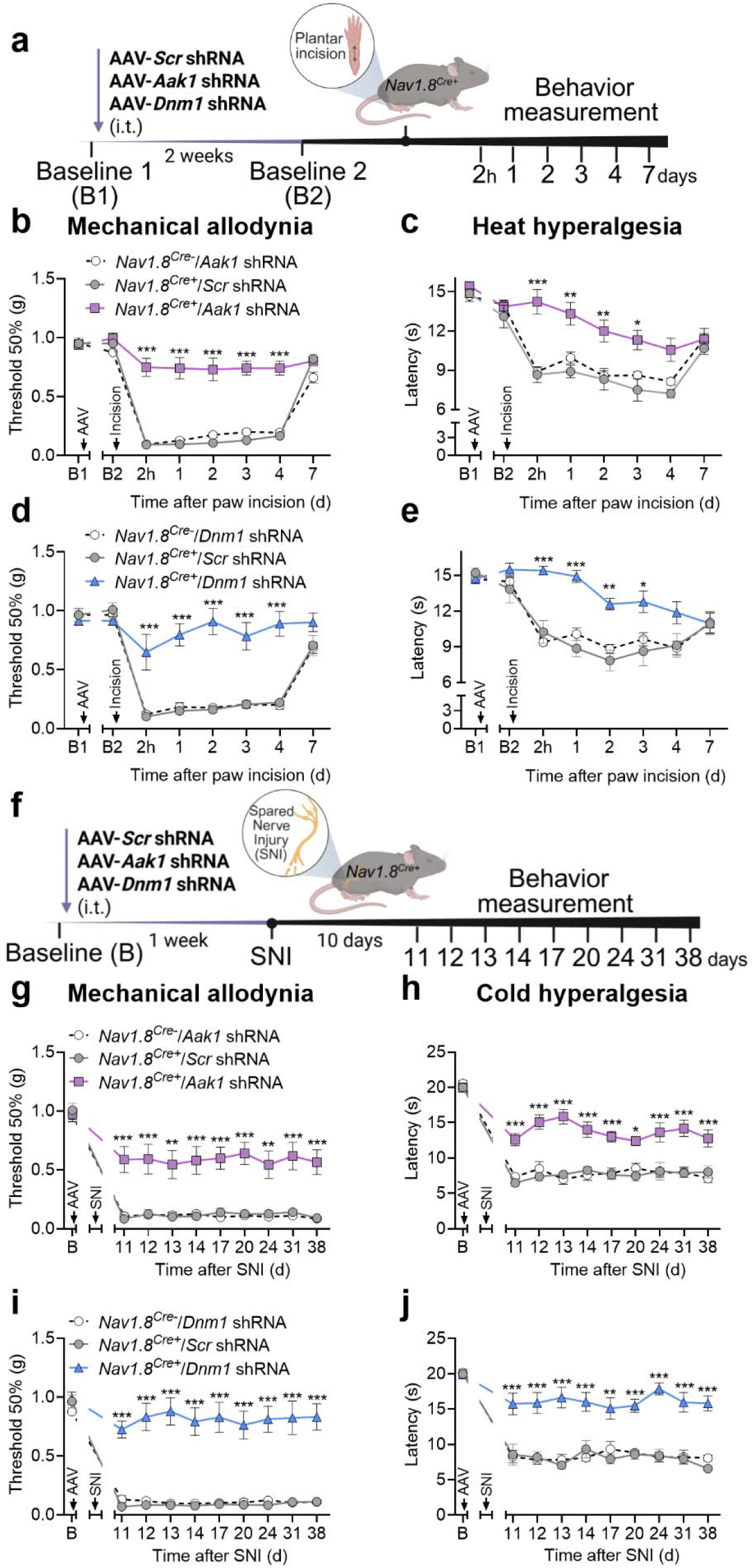
Selective knockdown of AAK1 or Dnm1 in nociceptors prevents postoperative and neuropathic pain. Experimental timeline of the postoperative pain model induced by plantar incision (a). Mechanical allodynia (b, d) and heat hyperalgesia (c, e) of the plantar incision in *Nav1.8^Cre^* male and female mice, measured from 2 hours to 7 days after incision. Mice received intrathecal (i.t.) injections of AAV *Aak1*, *Dnm1* or their respective *scrambled* controls (*Scr)* shRNA 2 weeks prior to the incision, n=7-8 mice per group. Experimental timeline of the neuropathic pain model induced by Spared Nerve Injury (SNI) (f). Mechanical allodynia (g, i) and cold hyperalgesia (h, j) in the ipsilateral paw of SNI *Nav1.8^Cre^* male and female mice, measured from 11-38 days post-surgery. Mice received i.t. injections of AAV *Aak1*, *Dnm1* or their respective *Scr* shRNA one week before the SNI surgery, n=6-8 mice per group. Data are presented as Mean ± SEM. *P<0.05, **P<0.01, ***P<0.001 vs. *Nav1.8^Cre-^*. Two-way ANOVA with Sídák multiple comparisons test. B1: baseline 1, pre-AAV injection; B2: baseline 2, two weeks post-AAV injection.

### Selective knockdown of AAK1 or Dnm1 in nociceptors prevents neuropathic pain

To investigate the effects of AAK1 and Dnm1 knockdown in a preclinical model of neuropathic pain, Nav1.8^Cre+^ or control (Nav1.8^Cre-^) mice underwent Spared Nerve Injury (SNI) surgery. AAV *Aak1*, *Dnm1* or *Scr* shRNA was administered by i.t. injection 1 week before the surgery. Mechanical allodynia and cold hyperalgesia of the operated (left, ipsilateral) and non-operated (right, contralateral) hindpaws were assessed using von Frey filaments and cold plate test, respectively, starting 10 days after SNI surgery, and monitored daily (**Fig. 2f**). SNI surgery induced robust and persistent mechanical allodynia and cold hyperalgesia for at least 38 days (**Fig. 2g-j**). AAV *Aak1* shRNA and AAV *Dnm1* shRNA sustainedly prevented SNI surgery-induced mechanical allodynia and cold hyperalgesia for the full 38 days compared to AAV *Scr* shRNA in male and female Nav1.8^Cre+^ mice, with no effects in Nav1.8^Cre-^mice. None of the treatments affected withdrawal responses of the contralateral (non-operated) paw to mechanical stimuli (**Fig. S3e, f**).

### Selective knockdown of AAK1 or Dnm1 in nociceptors preserves normal behavior and the protective reflexive response

Interference with normal behavior could impair the reflexive defensive responses that protect against injury and confound the assessment of nociception. To determine whether the selective knockdown of AAK1 or Dnm1 in nociceptors alters baseline behavior, AAV *Aak1*, *Dnm1* or *Scr* shRNA was injected i.t. in naïve mice. Two weeks after virus injection, AAK1 or Dnm1 knockdown did not affect baseline mechanical or heat sensitivity (**Fig. S4a-d**). Motor coordination, assessed by the rotarod test, was likewise unaffected, as latency to fall was similar across groups (**Fig. S4e**). Locomotor, exploratory and grooming behaviors were assessed using a behavioral spectrometer during 20-minute recordings. AAK1 or Dnm1 knockdown induced minor changes in grooming, visits to the center and still time behavior, but did not alter average velocity, track length, locomotory activity, ambulation or wall distance when compared with AAV *Scr* shRNA (**Fig. S4f-m**).

Together, these findings demonstrate that selective silencing AAK1 or Dnm1 in nociceptors produces robust and sustained suppression of postoperative and neuropathic pain without affecting baseline sensory or motor function.

### Selective knockdown of AAK1 and Dnm1 in nociceptors reduces post-synaptic current amplitude evoked by optogenetic activation of Nav1.8 afferent fibers

To investigate the contribution of AAK1 and Dnm1 in maintaining functional synaptic signaling from nociceptive afferent neurons to the spinal dorsal horn, we selectively activated Nav1.8+ve fibers *via* optogenetic stimulation of parasagittal spinal cord slices from mice injected with AAV *Aak1*, *Dnm1* or *Scr* shRNA (i.t.) at least 2 weeks prior. Expression of Cre-dependent AAV shRNA was targeted to Nav1.8+ve fibers that expressed light activated channel-rhodopsin-2 (ChR2). Optogenetically-evoked postsynaptic currents (oPSCs) recorded from superficial dorsal horn (lamina I-II) neurons in whole-cell voltage clamp configuration were significantly reduced by AAV *Aak1* shRNA (AAV *Scr* shRNA, 1.16 ± 0.29 nA; AAV *Aak1* shRNA, 0.55 ± 0.12 nA; p = 0.027, unpaired t-test, **Fig. 3b**) and by AAV *Dnm1* shRNA (AAV *Scr* shRNA, 1.47 ± 0.24 nA; *Dnm1* shRNA, 0.46 ± 0.15 nA; p = 0.0019, unpaired t-test, **Fig. 3c**). Repeating the stimuli every 10 seconds (0.1 Hz) showed no significant attenuation over 10 pulses, indicating that response to slow stimuli is maintained (P >0.5 compared to first pulse, mixed effects analysis with Dunnett’s multiple comparisons test, **Fig. S5a**).

**Figure 3.**
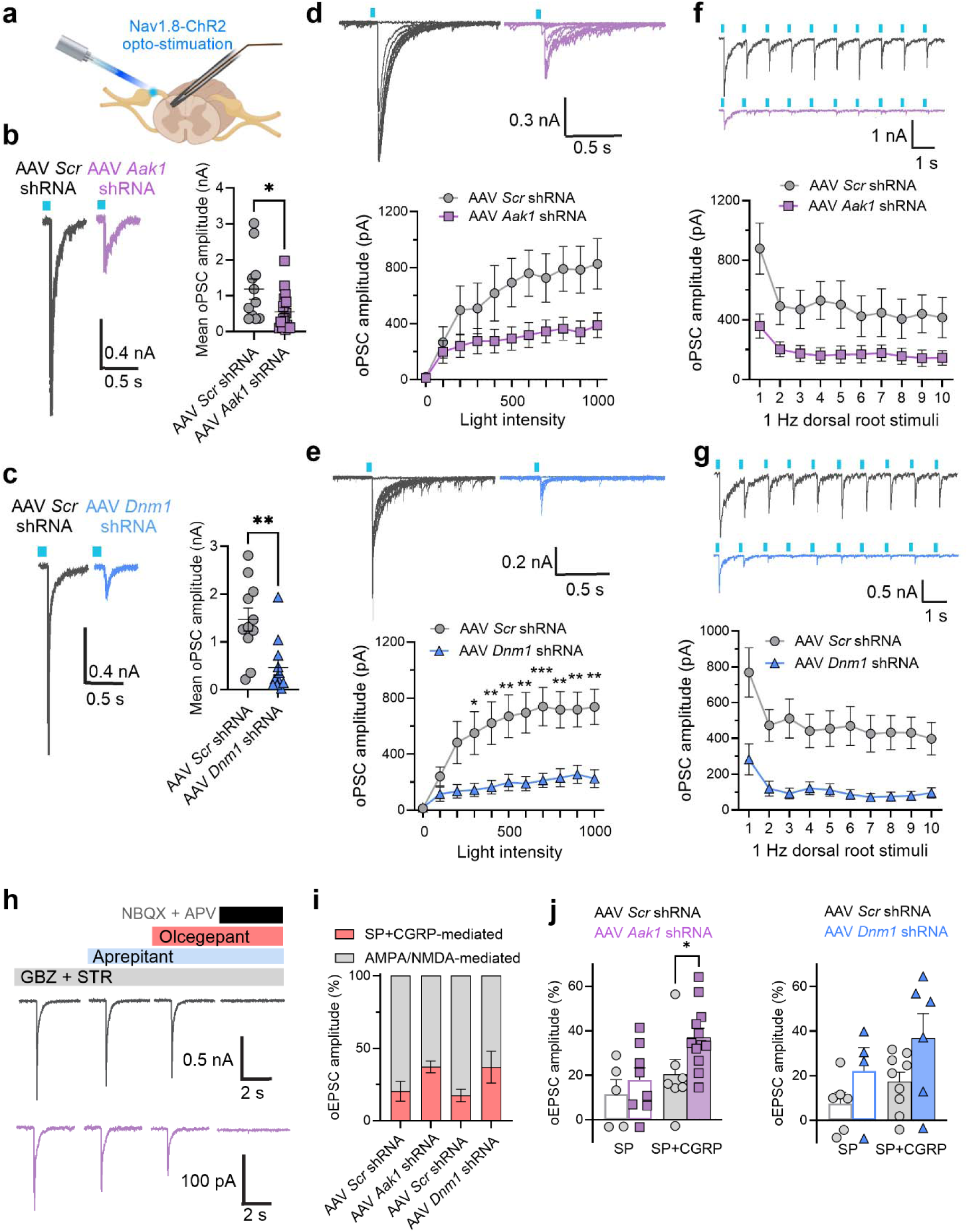
Nav1.8-afferent mediated synaptic signaling in the dorsal horn is decreased following selective knockdown of AAK1 and Dnm1. The dorsal root entry zone in parasagittal lumbar spinal cord slices from mice expressing ChR2 in Nav1.8 neurons was stimulated with a 465 nm light pulse (10ms) and optogenetically-evoked postsynaptic currents (oPSCs) recorded from dorsal horn neurons in the superficial laminae (a). Traces show oPSCs in response to a light pulse (blue square) in neurons from mice injected with AAV *Aak1* or *Dnm1* shRNA compared to their respective *scrambled* controls (*Scr)* shRNA (b, c). Traces and individual data points represent an average of 3 oPSCs per neuron (0.1 Hz). Representative traces and mean data of input-output responses of oPSCs from spinal cord slices in response to increasing light intensity in AAV *Aak1* and *Dnm1* shRNA compared to AAV *Scr* shRNA (d, e). Representative traces showing oPSC responses to 1Hz presynaptic stimulation in AAV *Aak1* and *Dnm1* shRNA compared to AAV *Scr* shRNA (f, g). Representative traces show oEPSC responses recorded in the presence of gabazine (GBZ) and strychnine (STR), that were sequentially blocked by antagonists targeting SP (aprepitant, 1 μM) and CGRP (olcegepant, 1 μM), followed by glutamatergic blockers NBQX and APV (h). Proportion of total oESPC (%) mediated by SP and CGRP, compared to glutamate in AAV *Aak1* and *Dnm1* shRNA-treated spinal cord neurons compared to controls (i). Comparison of neuropeptide contribution to total oESPC in AAV *Aak1* and *Dnm1* shRNA and control groups, showing SP-mediated and SP + CGRP-mediated components (j). Mean±SEM. *P<0.05, **P<0.01, ***P<0.001 versus control. Unpaired t-test (b, c, j) or Two-way ANOVA with Sídák multiple comparisons test (e).

To investigate postsynaptic responses over a range of stimulus intensities, the 465 nm LED light was increased in brightness from 0 (minimum) to 1000 (maximum) in increments of 100. The intensity required to elicit maximal synaptic responses was thereby determined. AAV *Aak1* or *Dnm1* shRNA and their respective AAV *Scr* shRNA-treated groups showed a gradual increase in oPSC as light intensity increased (**Fig. 3d, e**), with the AAV *Dnm1* shRNA showing significant differences in amplitude compared to the AAV *Scr* shRNA by the third increment of light intensity (2-way ANOVA, with Šídák’s multiple comparisons). When data were normalized to the maximal current, there were no significant difference in current amplitude between groups (P >0.5, mixed effects analysis with Tukey’s multiple comparisons), which suggests that the differences observed were due to changes in amplitude, rather than level of stimulation required to elicit a response.

To investigate whether postsynaptic currents in the AAV *Aak1* shRNA and AAV *Dnm1* shRNA groups were more likely to attenuate during 1 Hz dorsal root stimulation, current amplitudes in response to optogenetic activation were compared over 10 seconds. All groups showed a reduction in amplitude between the first and second stimuli (**Fig. 3f, g**), with a significant decrease in AAV *Aak1* shRNA and AAV *Dnm1* shRNA groups observed when data was normalized to the first pulse (**Fig. S5b**, p = 0.011 for AAV *Aak1* shRNA, p <0.0001 for AAV *Dnm1* shRNA, compared to AAV *Scr* shRNAs, 2way ANOVA), which suggests that current amplitude less sustained in AAV *Aak1* or *Dnm1* shRNA groups. Applying a pulse of 2 Hz showed a faster decline in amplitude with the AAV *Aak1* shRNA and AAV *Dnm1* shRNA groups (**Fig. S5c**) and confirmed that the elicited excitatory currents originated from C- and Aδ-fibers, which would be expected to fail at 2 Hz in a naïve (non-pain) state.

Since dorsal root-evoked glutamatergic currents are modulated by synaptically released neuropeptides(*22*), we sought to determine the contribution of SP and CGRP when expression of AAK1 or Dnm1 was downregulated. To determine the component of the total excitatory postsynaptic current mediated by SP and CGRP, the neurokinin 1 receptor (NK_1_R) antagonist aprepitant (1 μM), followed by aprepitant and the CGRP receptor antagonist olcegepant (1 μM), were applied sequentially during the recording (**Fig. 3h**). AAV *Aak1* shRNA resulted in an increase in the proportion of total current mediated by neuropeptides (**Fig. 3i**), with a significant increase in the percentage of SP + CGRP component (P = 0.037 compared to AAV *Scr* shRNA, unpaired t-test, **Fig. 3j**). Since the total current amplitude is decreased in AAV *Aak1* shRNA and AAV *Dnm1* shRNA treated groups, this result suggests that glutamate is preferentially reduced in the AAV *Aak1* shRNA treated group. To confirm that the remaining current was glutamatergic, the AMPA (NBQX) and NMDA (APV) antagonists were applied to the slice at the end of the recording. NBQX and APV abolished the remaining evoked current, confirming that residual synaptic transmission was primarily glutamatergic.

Together, these findings establish AAK1- and Dnm1-dependent endocytic mechanisms in nociceptors as essential drivers of neurotransmission and pain hypersensitivity and highlight their potential as targeted analgesic strategies.

### Development, characterization and targeting of LNPs encapsulating CRISPR/dCas9-R mRNA

To translate these findings into a therapeutic strategy, we designed an LNP-based CRISPR/dCas9 gene repression (dCas9-R) platform to achieve sustained and reversible gene silencing *in vivo*. LNPs analogous to the FDA-approved MC3-based *Onpattro* formulation(*17, 23*) were generated to encapsulate mRNA encoding CRISPR/dCas9-R together with synthetic sgRNAs targeting the promoter regions of the mouse *Aak1* or *Dnm1* genes (sequences in **Table S2**). The dCas9-R construct consists of endonuclease-dead Cas9 fused to ZIM3(*24*)-MeCP2(*25*) transcriptional and D3A-D3L(*26*) epigenetic repressor domains, enabling targeted suppression of gene expression without permanent genome editing. Such epigenetic repression is reversible and nonpermanent, features that are advantageous for therapeutic applications(*24–26*). All LNP-encapsulated CRISPR/dCas9-R formulations exhibited uniform particle size range of 70 and 90 nm, zeta potentials below 0.2 mV and encapsulation efficiencies of approximately 90 % or greater (**Table S3**).

To assess tissue targeting, LNP-encapsulated Cre recombinase mRNA (1µg/µl, 5µl) was delivered intrathecally to CAG^floxStop-tdTomato^ (Ai14) mice. The Ai14-Cre system is a widely used genetic model for evaluating gene editing e[ciency and tracing specific cell types *in vivo*(*27, 28*). Ai14 mice carry a *loxP*-flanked STOP cassette upstream of the tdTomato reporter, which is robustly expressed following Cre-mediated recombination (**Fig. 4a**). In mice treated with LNP-Cre mRNA, we observed widespread tdTomato expression throughout the DRG, with prominent accumulation in neurons, satellite glial cells, and macrophages after 5 days (**Fig. 4b**). In contrast, minimal expression was detected in the spinal cord and brain, primarily targeting the meninges (**Fig. 4c, d**). These findings indicate that intrathecally delivered LNPs primarily exhibit local distribution and transduction within the DRG.

**Figure 4.**
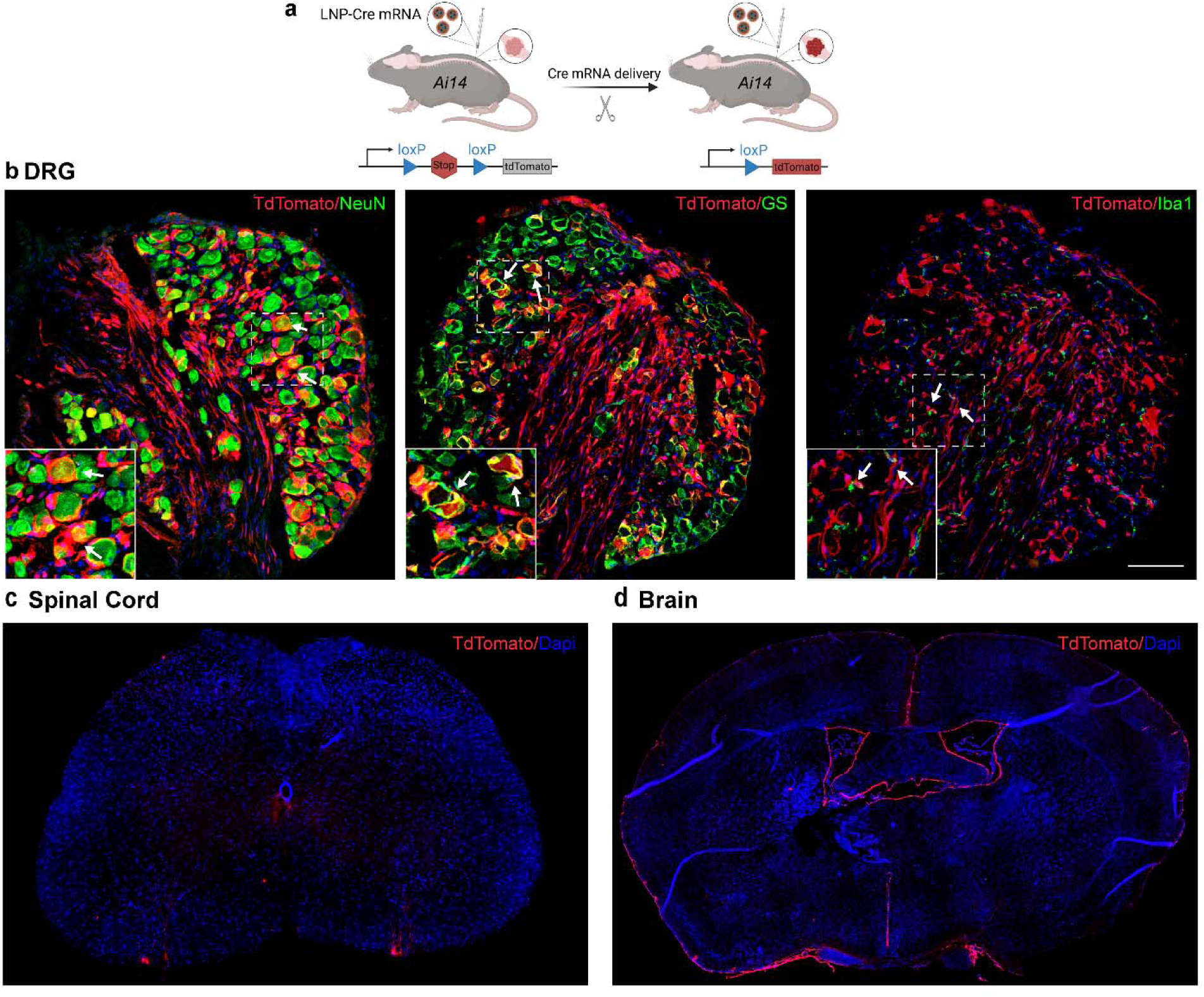
LNPs mediated Cre mRNA delivery via intrathecal injection in Ai14 mice. Schematic of LNP-mediated Cre mRNA delivery activating tdTomato expression in Ai14 mice (a). Representative DRG images showing tdTomato expression in neurons (NeuN), satellite glial cells (GS) and macrophages (Iba1) following intrathecal injection of LNP-Cre mRNA (1 µg/µl, 5 µl) (b). Representative spinal cord (c) and brain (d) images from injected mice. Scale bar: 100 µm.

### LNP-encapsulated CRISPR/dCas9-R mRNA targeting AAK1 or Dnm1 provides knockdown of targets in DRG

To validate target downregulation, LNP-encapsulated CRISPR/dCas9-R with a sgRNA targeting the promoter region of the mouse *Aak1* or *Dnm1* gene, or a non-targeting sgRNA negative control (NC, 300ng/ 5µl) was administered to mice (i.t., L4/L5). RNAScope revealed that LNP dCas9-R AAK1 or LNP dCas9-R Dnm1, but not LNP dCas9-R NC, reduced the expression of *Aak1* and *Dnm1* mRNA in DRG neurons by 45 ± 7% and 25 ± 8%, respectively, after 3 days compared with PBS (control, 5 µl) (**Fig. 5a, b**). Expression of *Aak1* and *Dnm1* mRNA in the spinal cord was unaffected (**Fig. S6a-d**).

**Figure 5.**
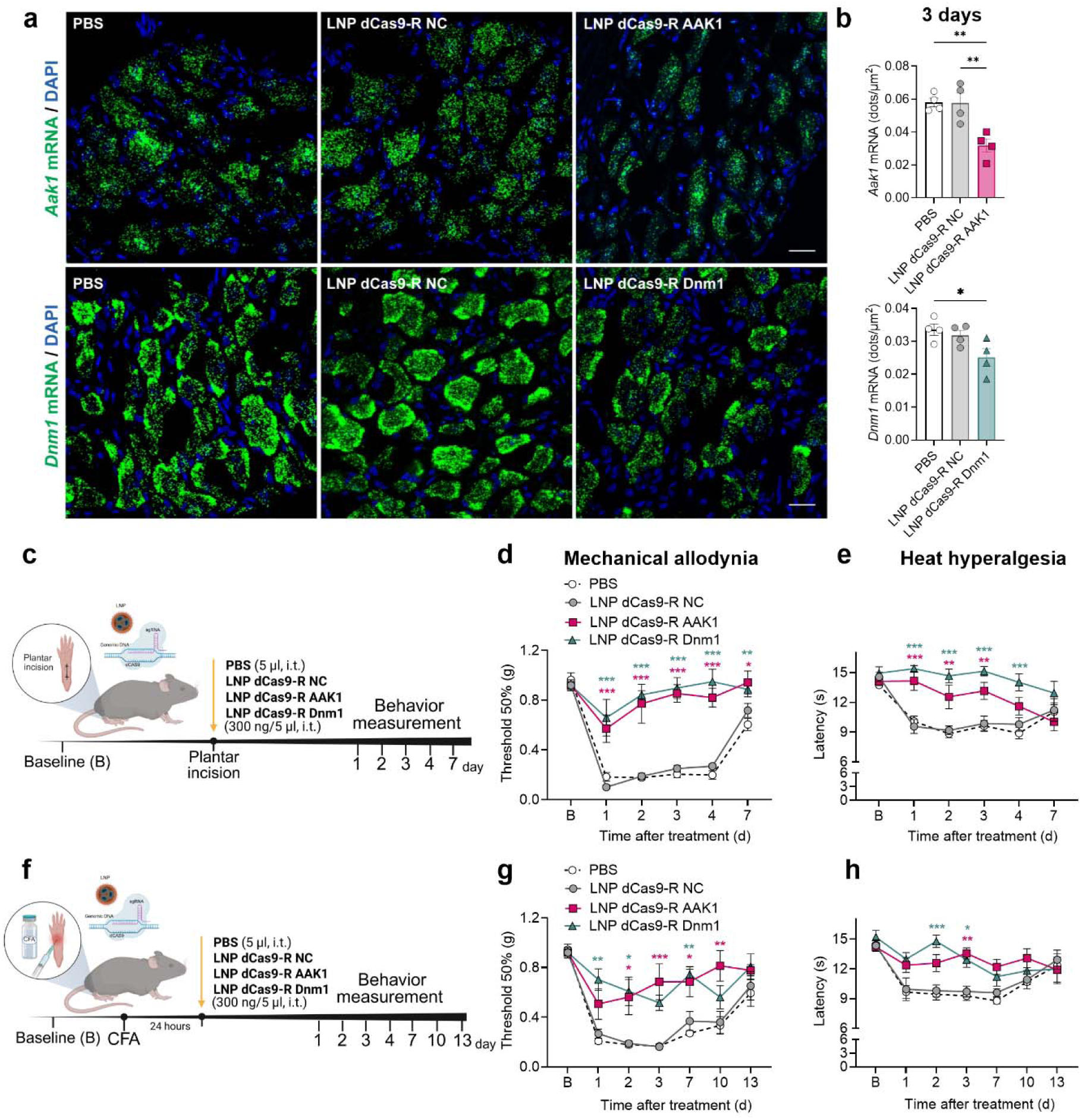
Effect of LNP-encapsulated CRISPR/dCas9-R AAK1 or Dnm1 on postoperative and inflammatory pain. RNAScope localization (a) and quantification (number of dots per area) (b) of *Aak1* and *Dnm1 mRNA* expression in mouse DRG at 3 days after intrathecal (i.t.) injection of LNP dCas9-R AAK1, LNP dCas9-R Dnm1, LNP dCas9-R NC (300 ng/ 5µl, i.t.) or PBS, n= 4 mice per group. Experimental timeline of postoperative pain model induced by plantar incision (c). Mechanical allodynia (d) and heat hyperalgesia (e) of the plantar incision mice measured 1-7 days after i.t. injection of LNP dCas9-R AAK1, LNP dCas9-R Dnm1, LNP dCas9-R NC (300 ng/ 5µl, i.t.) or PBS, n=6 mice per group. Experimental timeline of inflammatory pain model induced by intraplantar injection of Complete Freund’s Adjuvant (CFA) (f). Mechanical allodynia (g) and heat hyperalgesia (h) of CFA mice measured 1-13 days after i.t. injection of LNP dCas9-R AAK1, LNP dCas9-R Dnm1, LNP dCas9-R NC (300 ng/ 5µl, i.t.) or PBS, n=6 mice per group. Data are presented as Mean±SEM. *P<0.05, **P<0.01, **p<0.001 vs. PBS or LNP dCas9-R NC. 2-way ANOVA, Sídák multiple comparisons test or 1-way ANOVA, Tukey multiple comparisons test. NC, negative control.

To determine whether LNP-encapsulated CRISPR/dCas9-R mRNA gene editing could offer a new therapeutic approach to pain treatment, we investigated the effects of LNP dCas9-R AAK1 or LNP dCas9-R Dnm1 in preclinical models of postoperative, inflammatory, neuropathic and osteoarthritis pain in mice.

### LNP-encapsulated CRISPR/dCas9-R mRNA targeting AAK1 or Dnm1 prevents postoperative pain and reverses inflammatory pain

Postoperative pain was induced by plantar incision and evaluated from 1 to 7 days after incision (**Fig. 5c**). LNP dCas9-R AAK1, LNP dCas9-R Dnm1, LNP dCas9-R NC (300ng/ 5µl, i.t.) or PBS was administered by i.t. injection immediately after the incision. LNP dCas9-R AAK1 and LNP dCas9-R Dnm1 caused a nearly complete and long-lasting inhibition of incision-evoked mechanical allodynia and heat hyperalgesia for at least 4 days when compared to PBS (**Fig. 5d, e**). LNP dCas9-R AAK1 prevented mechanical allodynia by 93 ± 7% of baseline at day 3 and heat hyperalgesia by 92 ± 21% of baseline at day 1. LNP dCas9-R Dnm1 prevented mechanical allodynia and heat hyperalgesia by 100% of baseline at day 4. The LNP dCas9-R NC had no effect.

Inflammatory pain was induced by intraplantar injection of Complete Freund’s Adjuvant (CFA) or vehicle (control) into the hindpaw. LNP dCas9-R AAK1, LNP dCas9-R Dnm1, LNP dCas9-R NC (300ng/ 5µl, i.t.) or PBS was administered by i.t. injection 24 h after CFA, and mechanical allodynia and heat hyperalgesia were assessed daily (**Fig. 5f**). CFA-induced mechanical allodynia and heat hyperalgesia for at least 13 days (**Fig. 5g, h**). LNP dCas9-R AAK1 and LNP dCas9-R Dnm1 caused a long-lasting inhibition of mechanical allodynia for 10 and 7 days, respectively, when compared to PBS (**Fig. 5g**). LNP dCas9-R AAK1 and LNP dCas9-R Dnm1 also reversed thermal hyperalgesia for 3 days after treatment (**Fig. 5h**). LNP dCas9-R AAK1 reversed mechanical allodynia by 74 ± 16% of baseline and heat hyperalgesia by 88 ± 12% of baseline at day 3. LNP dCas9-R Dnm1 reversed mechanical allodynia by 81 ± 4% of baseline at day 7 and heat hyperalgesia by 100% of baseline at day 2. The LNP dCas9-R NC had no effect.

None of the treatments (plantar incision, intraplantar CFA, i.t. LNPs) affected withdrawal responses of the contralateral (non-incision/non-injected) paw to mechanical stimuli (**Fig. S7a, b**).

### LNP-encapsulated CRISPR/dCas9-R mRNA targeting AAK1 or Dnm1 reverses neuropathic and osteoarthritis pain and provides sustained knockdown of targets in DRG

Neuropathic pain was induced by SNI surgery of the hindpaw. LNP dCas9-R AAK1, LNP dCas9-R Dnm1, LNP dCas9-R NC (1µg/ 5µl, i.t.) or PBS was administered i.t. 10 days after surgery and mechanical allodynia and cold hyperalgesia were assessed daily (**Fig. 6a**). LNP dCas9-R AAK1 and LNP dCas9-R Dnm1 caused a long-lasting inhibition of SNI-induced mechanical allodynia and cold hyperalgesia for 21 and 14 days, respectively, when compared to PBS (**Fig. 6b, c**). LNP dCas9-R AAK1 reversed mechanical allodynia by 54 ± 6% of baseline at day 7 and heat hyperalgesia by 66 ± 15% of baseline at day 4. LNP dCas9-R Dnm1 reversed mechanical allodynia by 63 ± 10% of baseline and heat hyperalgesia by 77 ± 8% of baseline at day 4. The LNP dCas9-R NC had no effect.

**Figure 6.**
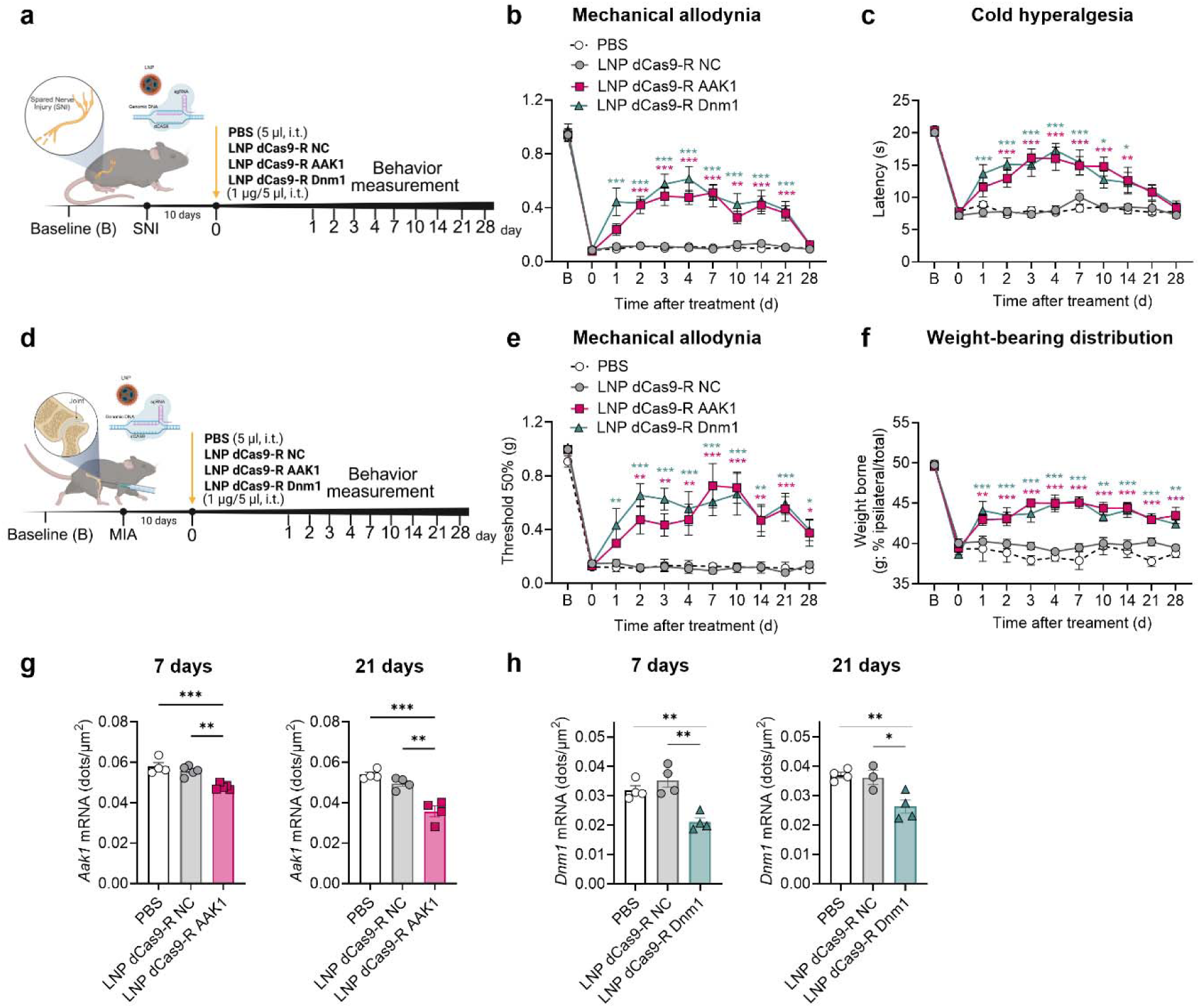
Effect of LNP-encapsulated CRISPR/dCas9-R AAK1 or Dnm1 on neuropathic and osteoarthritis pain. Experimental timeline of neuropathic pain model induced by Spared Nerve Injury (SNI) (a). Mechanical allodynia (b) and cold hyperalgesia (c) in the ipsilateral paw of SNI mice, measured 1-28 days after intrathecal (i.t.) injection of LNP dCas9-R AAK1, LNP dCas9-R Dnm1, LNP dCas9-R NC (1 µg/ 5µl, i.t.) or PBS, n=8 mice per group. Experimental timeline of osteoarthritis pain model induced by intra-articular injection of monosodium iodoacetate (MIA) (d). Mechanical allodynia in the ipsilateral paw (e) and weight bearing distribution across the 2 hindlimbs (f) of MIA mice, measured 1-28 days after i.t. injection of LNP dCas9-R AAK1, LNP dCas9-R Dnm1, LNP Cas9-R NC (1 µg/ 5µl, i.t.) or PBS, n=6 mice per group. RNAScope quantification (number of dots per area) of *Aak1* (g) and *Dnm1* (h) *mRNA* expression in mouse DRG at 7 and 21 days after i.t. injection of LNP dCas9-R AAK1, LNP dCas9-R Dnm1, LNP dCas9-R NC (1 µg/ 5µl, i.t.) or PBS, n= 4 mice per group. Data are presented as Mean±SEM. *P<0.05, **P<0.01, ***P<0.001 vs. PBS or LNP dCas9-R NC. 2-way ANOVA, Sídák multiple comparisons test or 1-way ANOVA, Tukey multiple comparisons test. NC, negative control.

We then investigated the effects of LNP dCas9-R AAK1 or LNP dCas9-R Dnm1 in a clinically relevant model of chronic musculoskeletal pain. Osteoarthritis pain was induced by intra-articular injections of monosodium iodoacetate (MIA) (1 mg/mL) via the infra-patellar ligament of the left knee. LNP dCas9-R AAK1, LNP dCas9-R Dnm1, LNP dCas9-R NC (1µg/ 5µl, i.t.) or PBS was administered i.t. 10 days after MIA and mechanical allodynia and weight-bearing distribution were assessed daily (**Fig. 6d**). MIA induced sustained mechanical allodynia and weight-bearing asymmetry. LNP dCas9-R AAK1 and LNP dCas9-R Dnm1 caused prolonged inhibition of MIA-induced mechanical allodynia and ameliorated weight-bearing symmetry for at least 28 days when compared to PBS (**Fig. 6e, f**). LNP dCas9-R AAK1 reversed mechanical allodynia by 73 ± 14% of baseline and weight-bearing symmetry by 64 ± 4% of baseline at day 7. LNP dCas9-R Dnm1 reversed mechanical allodynia by 66 ± 12% of baseline at day 10 and weight-bearing symmetry by 66 ± 2% of baseline at day 7. The LNP dCas9-R NC had no effect.

None of the treatments (SNI surgery, intra-articular MIA, i.t. LNPs) affected withdrawal responses of the contralateral (non-operated/non-injected) paw to mechanical stimuli (**Fig. S7c, d**).

LNP-encapsulated CRISPR/dCas9-R mRNA (1µg/ 5µl, i.t.) induces sustained knockdown of the targeted genes. LNP dCas9-R AAK1 and LNP dCas9-R Dnm1, but not LNP dCas9-R NC, reduced *Aak1* and *Dnm1* mRNA levels in DRG neurons. *Aak1* mRNA expression was reduced by 17 ± 1% and 34 ± 5% after 7 and 21 days, respectively, compared to PBS (**Fig. 6g**). *Dnm1* mRNA expression was reduced by 32 ± 5% and 29 ± 6% after 7 and 21 days, respectively, compared to PBS (**Fig. 6h**). Expression of *Aak1* and *Dnm1* mRNA in the spinal cord was unaffected (**Fig. S6a-d**).

### LNP-encapsulated CRISPR/dCas9-R mRNA targeting AAK1 or Dnm1 does not affect sensitivity, locomotor, exploratory and grooming behavior in naïve mice

To assess the potential short and long-term effects of the CRISPR/dCas9-R mRNA-encapsulated LNPs on sensitivity, motor coordination and non-evoked behavior, LNP dCas9-R AAK1, LNP dCas9-R Dnm1, LNP dCas9-R NC (300 ng and 1µg/ 5µl) or PBS was injected i.t. in naïve mice. None of the treatments affected mechanical, heat or cold sensitivity when assessed at 2 days (300 ng/ 5µl, i.t.) or at 3, 7, 14 and 21 days (1µg/ 5µl, i.t.) after treatment (**Fig. 7a-c, e-g**). Motor coordination was likewise unaffected, as latency to fall was similar across groups (**Fig. 7d, h**). Locomotor, exploratory and grooming behaviors were recorded for 20 minutes on day 2 (300 ng/ 5µl, i.t.) or days 7 and 21 (1µg/ 5µl, i.t.) after the treatment. The lower dose of LNP dCas9-R AAK1, LNP dCas9-R Dnm1 or LNP dCas9-R NC did not affect average velocity, track length, locomotor activity, ambulation, grooming, wall distance, visits to the center and still time parameters when compared to PBS (**Fig. 7i-p, Day 2**). The higher dose of LNP dCas9-R AAK1 and LNP dCas9-R Dnm1 caused a small but significant increase in average velocity and track length, without affecting the other parameters at day 7 when compared to LNP dCas9-R NC or PBS (**Fig. 7i-p, Day 7**). However, no behavioral alterations were observed on day 21 (**Fig. 7i-p, Day 21**). Thus, LNP-encapsulated CRISPR/dCas9-R mRNA targeting AAK1 or Dnm1 causes *Aak1* and *Dnm1* mRNA downregulation without discernible effects on sensitivity, locomotor activity and non-evoked behavior in naïve mice.

**Figure 7.**
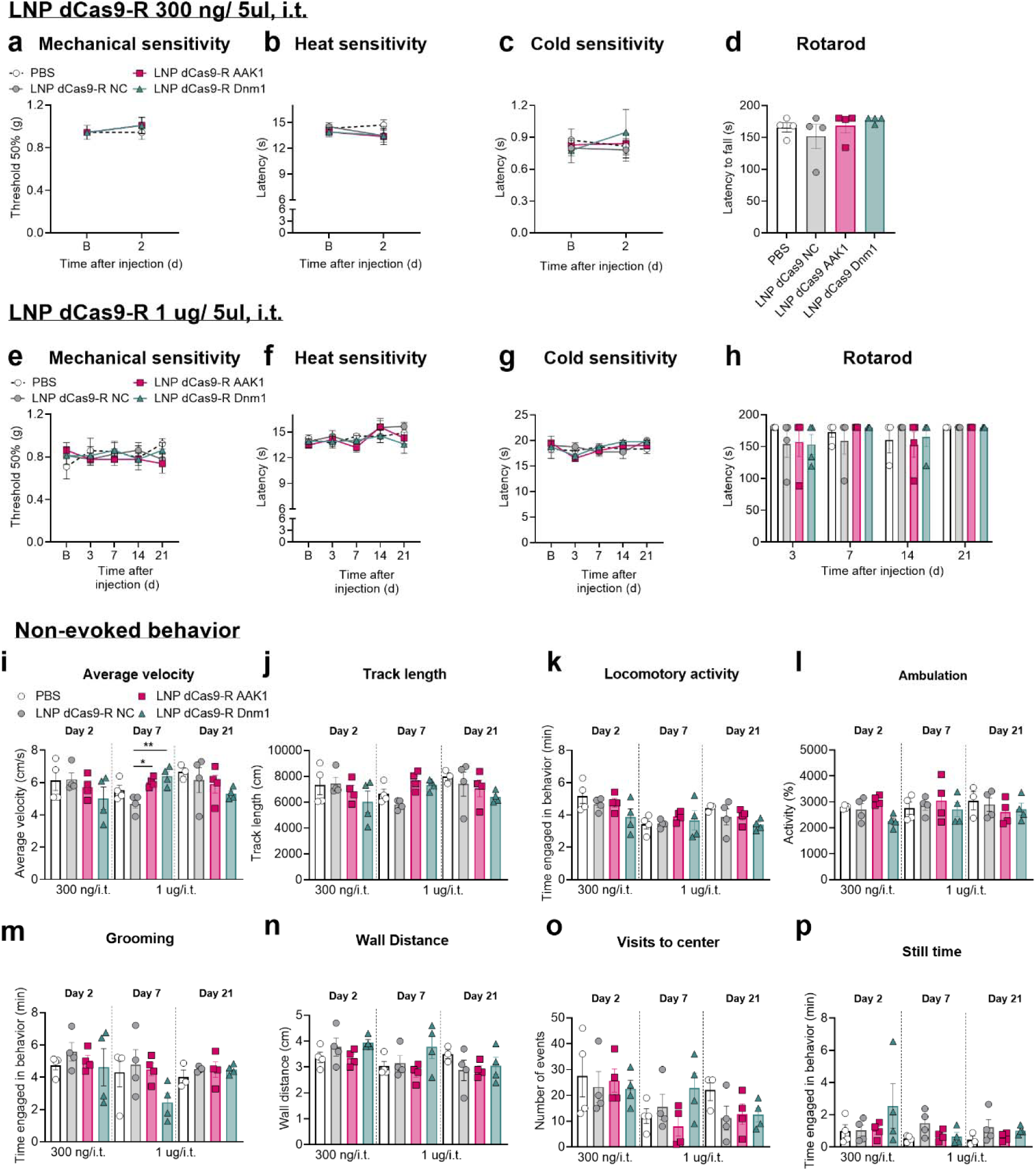
LNP dCas9-R AAK1 or Dnm1 mRNA does not affect sensitivity, locomotor, exploratory and grooming behavior in naïve mice. Mechanical (a), heat (b) and cold sensitivity (c), and locomotor activity (rotarod) (d) in naïve mice 2 days after intrathecal (i.t.) administration of a low dose of LNP dCas9-R AAK1, LNP dCas9-R Dnm1, LNP dCas9-R NC (300 ng/ 5µl, i.t.) or PBS, n=4 mice per group. Mechanical (e), heat (f) and cold sensitivity (g), and locomotor activity (rotarod) (h) in naïve mice 3, 7, 14 and 21 days after i.t. injection of a higher dose of LNP dCas9-R AAK1, LNP dCas9-R Dnm1, LNP dCas9-R NC (1 ug/ 5µl, i.t.) or PBS, n=4 mice per group. (i-p) Non-evoked behavior recorded for 20 min in naïve mice at days 2 (300 ng/ 5µl), 7 or 21 (1µg/ 5µl) after i.t injection of LNP dCas9-R AAK1, LNP dCas9-R Dnm1, LNP dCas9-R NC or PBS, n=4 mice per group. Data are presented as Mean±SEM. *P<0.05, **P<0.01 vs. LNP dCas9-R NC or PBS. 1-way ANOVA, Tukey multiple comparisons test.

### Safety profile of LNP-encapsulated CRISPR/dCas9-R mRNA in mice

LNPs are the primary platform for RNA delivery but are frequently associated with inflammatory responses(*19, 29*). To assess the safety profile of intrathecally delivered LNP-encapsulated CRISPR/dCas9-R mRNA, we evaluated neuroinflammatory and systemic immune responses. Cytokine and chemokine levels were measured using a proteome profiler array (40 analytes) in DRG and spinal cord, and by Luminex® xMAP® technology (36 cytokines, chemokines and growth factors) in serum. LNP dCas9-R NC was delivered intrathecally, tissues and serum were collected at 6 h or 3 days post-injection. At 6 h post-injection, DRGs and spinal cord exhibited transient increases in pro-inflammatory cytokines, including Il-16, Il-1β, Timp-1, as well as chemokines such as CXCL10, CCL2, CXCL9 and SDF-1. By 3 days post-injection, all measured biomarkers had returned to baseline levels of PBS (**Fig. 8a, b**), coinciding with the time point at which LNP-encapsulated CRISPR dCas9-R mRNA produced an efficient antinociceptive effect. A similar transient inflammatory profile was observed in serum, with elevated inflammatory biomarkers at 6 h that resolved to baseline levels of PBS by 3 days post-injection (**Fig. 8c**).

**Figure 8.**
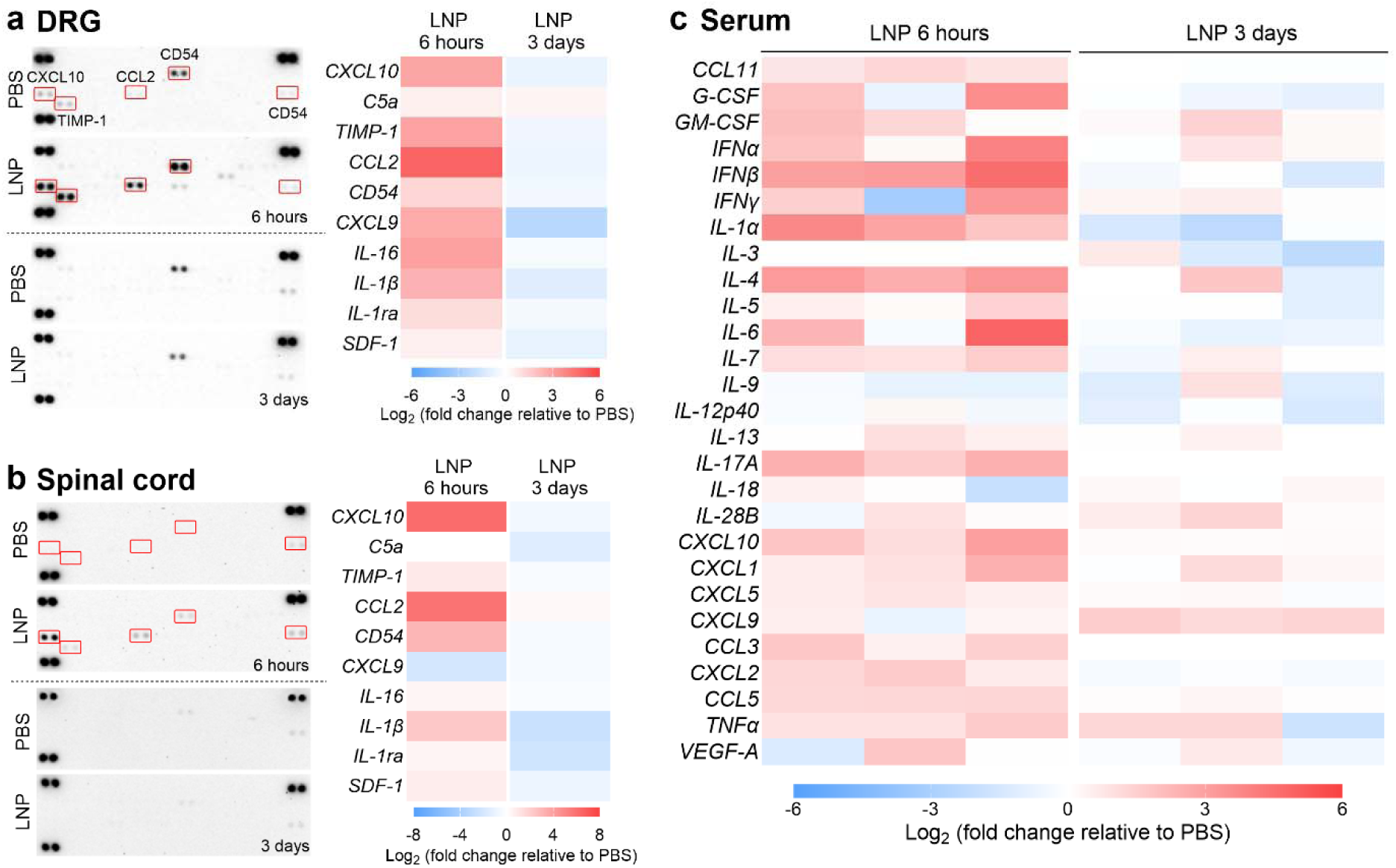
Safety profiles of LNP-mRNA for intrathecal injection. The expression levels of cytokines and chemokines in the DRG (a), spinal cord (b) and serum (c) of mice 6 hours or 3 days after intrathecal injection with LNP dCas9-R NC (1µg/ 5µl) or PBS, shown in blot array and heatmaps. DRG and spinal cord results indicate a pool of 5 mice. Serum results indicate three replicates. NC, negative control.

## Discussion

Our results show that SV endocytosis in nociceptors is a critical mechanism for sustained pain neurotransmission and that targeted disruption of this process provides effective and long-lasting pain relief. Using intrathecal AAV delivery of shRNA in Nav1.8^Cre^ mice, we first confirmed that the endocytic proteins AAK1 and Dnm1, selectively in nociceptors, are required to maintain synaptic transmission in nociceptive circuits and pain hypersensitivity. As a new therapeutic approach targeting these proteins, we developed an LNP-based CRISPR/dCas9-R delivery system to locally downregulate *Aak1* and *Dnm1* expression in the DRG, achieving broad and sustained pain relief across multiple preclinical pain models. Together, these findings elucidate a new mechanism of pain transmission and support a translational, gene modulation-based approach for non-opioid pain therapy.

### The requirement of endocytosis for pain transmission

A central unresolved question in our previous studies targeting endocytic mediators was the specific role of these mediators within nociceptors. Although genetic and pharmacological inhibition of key endocytic proteins, including AAK1 and Dnm1, suppresses synaptic transmission in nociceptive spinal circuits and attenuates pain-like behavior(*10, 11*), these approaches did not establish cell-type specificity. By selectively silencing *Aak1* or *Dnm1* in Nav1.8+ve neurons, we demonstrated that these endocytic proteins act within nociceptors to sustain neurotransmission and pain hypersensitivity. Nociceptor-specific *Aak1* or *Dnm1* knockdown markedly inhibited postoperative and neuropathic pain, without affecting normal sensitivity, locomotor or spontaneous behavior. Electrophysiological studies using specific optogenetic stimulation of nociceptors revealed a major role for AAK1 and Dnm1 in synaptic transmission within nociceptors and dorsal horn neurons. Selective AAK1 or Dnm1 knockdown in Nav1.8+ve afferents significantly reduced oPSCs in dorsal horn neurons, implicating a presynaptic mechanism due to a reduction in the probability of neurotransmitter release, consistent with limited vesicle availability. While responses were preserved at low-frequency stimulation (0.1 Hz), modest increases in frequency (1–2 Hz) induced pronounced synaptic depression following AAK1 or Dnm1 knockdown. These findings indicate that AAK1- and Dnm1-dependent endocytosis is required to sustain transmission during ongoing activity, a condition that more closely reflects pathological pain states. Disruption of SV endocytosis also altered the composition of synaptic signaling. Pharmacological blockage of NK_1_R and CGRP receptors revealed an increased proportional contribution of SP and CGRP to the total postsynaptic response following AAK1 knockdown. Given the overall reduction in current amplitude, this likely reflects preferential impairment of fast glutamatergic transmission rather than enhanced neuropeptide release. Thus, behavioral and electrophysiological data establish AAK1 and Dnm1 as necessary for presynaptic endocytic regulation in nociceptors, required for maintaining sustained pain signaling while preserving basal neurotransmission.

### Endocytosis inhibitors for pain relief

The identification of endocytic proteins as intrinsic regulators of nociceptors provides critical mechanistic insight into pain transmission, with implications for therapy. Our findings support a model in which AAK1- and Dnm1-dependent endocytosis replenishes the releasable SV pool required for sustained neurotransmitter release at nociceptor terminals. Disrupting this machinery likely depletes the SV pool and diminishes the release of excitatory neurotransmitters that drive second-order spinal neurons. Unlike strategies that target individual receptors or channels, interfering with SV recycling suppresses the common synaptic output of diverse nociceptive inputs. This convergence likely underlies the broad analgesic efficacy observed across postoperative and neuropathic pain models. Notably, partial disruption of SV endocytosis was sufficient to attenuate pain-like behaviors without impairing baseline sensory or motor function. This suggests that SV recycling is particularly critical during sustained nociceptor activity, making presynaptic endocytic regulation an appealing approach for reducing chronic pain while preserving essential protective sensation.

Nociceptor-specific genetic knockdown provided essential mechanistic validation and established a strong foundation for translational development. Gene therapy has proven to be particularly attractive for pain treatment. Transcription repression strategies, including CRISPR/dCas9-R-based systems, have demonstrated antinociceptive effects in inflammatory and neuropathic pain models(*12, 30, 31*). Building on our identification of endocytic proteins as regulators of nociceptive synaptic transmission, we developed an mRNA therapy strategy targeting this presynaptic mechanism. Local LNP-mediated delivery of mRNA encoding CRISPR/dCas9-R to silence either AAK1 or Dnm1 not only prevented the development of postoperative pain but also reversed established inflammatory pain and chronic neuropathic and osteoarthritis pain. Clinically, targeting the endocytic pathway is gaining traction, with AAK1 inhibitor LX9211 demonstrating promising efficacy and safety in phase II clinical trials for postherpetic neuralgia and diabetic neuropathy(*32, 33*). Notably, the long-lasting antinociceptive effect achieved with LNP-mediated CRISPR/dCas9-R distinguishes this approach from conventional pharmacological therapies that require repeated dosing, offering a potential advantage particularly in chronic pain treatment.

The therapeutic effects of LNP CRISPR/dCas9-R-mediated gene modulation closely mirrored those observed with nociceptor-specific knockdown, supporting the conclusion that this approach effectively targets the same presynaptic mechanisms. The local intrathecal delivery achieved efficient passive targeting of DRG neurons, as evidenced by the LNP biodistribution sites and the specific downregulation of DRG but not spinal cord neurons, while limiting the spread to the central nervous system. A previous study also reported that intrathecal administration of LNP containing mRNA successfully delivered to the DRGs(*34*). Notably, LNP-treated animals showed preserved baseline reflexes, locomotor and exploratory behavior, indicating that the treatment selectively attenuates pain hypersensitivity rather than causing global sensory or motor impairment. Importantly, the use of CRISPR/dCas9-R inhibits transcription without altering the DNA sequence in the genome and enables sustained yet reversible gene downregulation, providing a favorable safety profile by avoiding permanent genomic modification(*30, 35*). These findings underscore the translational promise of LNP CRISPR/dCas9-R as a platform for modulating pain-relevant genes in the peripheral nervous system.

### Limitations

Despite these promising results, some limitations warrant consideration. Although repression of *Aak1* or *Dnm1* produced robust and sustained analgesia, the potential for compensatory changes in other components of the SV trafficking machinery remains to be determined. Future studies will be required to assess whether repeated administration, and consequently prolonged gene repression, induces adaptive molecular or functional alterations that could influence efficacy or durability of analgesia. In addition, while local intrathecal delivery of LNP-CRISPR/dCas9-R was effective in the present study, the pharmacokinetics, safety and tolerability of repeated spinal dosing require further investigation. Although no major neuroinflammatory changes were detected under the conditions tested, more comprehensive analyses are needed to evaluate potential inflammatory responses, neurotoxicity and effects on neuronal integrity, particularly following long-term or repeated treatment. Finally, as with all CRISPR-based transcriptional repression strategies, the possibility of off-target gene regulation cannot be fully excluded and will require systematic transcriptomic and functional validation to establish specificity and safety as this approach advances toward clinical translation.

## Materials and Methods

### Study Design

The primary objective of this study was to investigate the role of SV endocytosis in nociceptors in driving chronic pain and to evaluate the therapeutic efficacy of a non-viral, LNP-mediated epigenetic silencing platform. We employed a multi-step approach beginning with target validation using AAV vectors to selectively knockdown AAK1 or Dnm1 in Nav1.8+ve nociceptors in Nav1.8^Cre+^ mice. Subsequently, we developed a translational platform using LNPs to deliver CRISPR/dCas9-R mRNA for reversible epigenetic repression of the same targets. Efficacy was evaluated across multiple preclinical models of pain (postoperative, inflammatory, neuropathic and osteoarthritis), utilizing evoked and non-evoked behavioral assessments. To evaluate synaptic transmission within spinal circuits, we used an electrophysiological approach combined with optogenetics in Nav1.8ChR2 mice to selectively activate nociceptor terminals.

Sample sizes of n=6-8 mice for behavior experiments were determined based on previous studies with these specific pain models to ensure sufficient statistical power. To ensure reproducibility, experiments were performed in at least two independent cohorts. Mice were randomly assigned to treatment or control groups using a web-based randomization procedure (http://www.randomizer.org/). All behavioral assessments and molecular analyses were performed by investigators blinded to the genetic background and treatment allocation of the animals. No animals were excluded from the study analysis. All experiments and procedures were approved by the New York University Institutional Animal Care and Use Committee and the Monash University Animal Ethics Committee. They were conducted following the guidelines established by the National Institute of Health, the International Association for the Study of Pain, and the National Center for the Replacement, Refinement, and Reduction of Animals in Research ARRIVE guidelines.

### AAV Formulation

#### Plasmid constructs

To obtain plasmids coding for short hairpin RNA targeting *mAak1*, *mDnm1* or *scrambled* (*Scr*) shRNA as negative control, the plasmid pAAV-CMV-LL-rev(GFP-5’ miR-30E-BfuAI-ORF_44bp BfuAI-3’ miR-30E)-rev(LL)-WPRE was obtained by Vectorbuilder. shRNAs targeting the gene of interest (*mAak1* P1/P2 or *mDnm1* P3/P4, Broad Institute, Genetic Perturbation Platform) or *Scr* shRNA (*Scr mAak1* P5/P6 and *mDnm1* P7/P8, Invivogen) were cloned by replacing the 44bp ORF with preannealed and phosphorylated oligonucleotides by using BfuAI compatible bases. Plasmid constructs were confirmed by Sanger sequencing. Primer sequences are available in **Table S1**.

#### AAVpro293T cell line

AAVpro 293T cells (#632273, Takara), were maintained in DMEM high glucose supplemented with 10% heat inactivated FBS, 4 mM L-glutamine, 1 mM penicillin/streptomycin and 1 mM sodium pyruvate at 37 °C in 5% CO_2_ and 95% O_2_. The day before transfection, cells were plated in DMEM supplemented with 2% tetracycline-free FBS.

#### AAV generation

Recombinant AAV particles (rAAVs) were produced by using triple transfection strategy as described previously(*36*). In brief, AAVpro 293T cells (#632273, Takara), were transfected with polyethylenimine (#23966, PEI, Polyscience) with a DNA:PEI ratio of 1:3. To obtain rAAVs, AAVpro 293T cells were transiently transfected with 2.5 mg total DNA (plasmid expressing genes of interest, pAdDeltaF6; #11287, Addgene and Rep/Cap, 1:1:1 molar ratio). To infect nociceptors with high efficiency as previously reported(*37*), Rep/Cap PHP.S was used (pUCmini-iCAP-PHP.S #103006, Addgene). rAAVs were extracted and isolated 72 h post-transfection, purified by iodixanol gradient ultracentrifugation, concentrated, and titrated using an RTqPCR assay (#6233 AAVpro Titration Kit, Takara) according to the manufacturer’s instructions.

### LNP CRISPR dCas9-R Formulation

#### sgRNA Design

sgRNA were designed using CRISPick (https://portals.broadinstitute.org/gppx/crispick/public). For dCas9-R experiments targeting AAK1 and Dnm1, three sgRNA were designed for each target, with the targeted locus falling within 0 – 250 bp downstream of the transcription start site (TSS) of the selected gene. A non-targeting sgRNA was designed as a negative control. Designed sgRNA were ordered as chemically synthesized sgRNA with end-modifications (Synthego, California, United States). Highly modified sgRNA was designed according to the motif described in Finn *et al.*(*38*)(**Table S2**)

#### mRNA Synthesis

Linear DNA templates for *in vitro* transcription containing a T7 promoter, 5’ and 3’ untranslated regions (UTRs), coding sequence and poly A tail were generated using PCR. All mRNA constructs were synthesized according to the HiScribe™ T7 mRNA Kit with CleanCap® Reagent AG protocol (New England Biolabs, Massachusetts, United States), using N1-methylpseudouridine-5’-triphosphate (TriLink BioTechnologies, California, United States) instead of uridine. The linear DNA template was removed by DNAse I treatment at 3717C for 30 minutes. The mRNA was precipitated using LiCl treatment and then stored in 1 mM sodium citrate solution, pH 6.4 at -8017C until use.

#### LNP Formulation

A lipid stock solution consisting of DLin-MC3-DMA (DC Chemicals, Shanghai, China), DSPC (Avanti Polar Lipids, Alabama, United States), cholesterol (Sigma-Aldrich, Massachusetts, United States), and DMG-PEG2000 (Avanti Polar Lipids, Alabama, United States) was prepared in ethanol at a molar ratio of 50:10:38.5:1.5 and a total lipid concentration of 20 mM.

For formulation, lipids were combined with mRNA (for biodistribution experiments) or dCas9-R mRNA and sgRNA (1:1, w/w) in 30 mM acetate buffer, pH 4.0 at a 1:3 lipid:RNA (v/v ratio, 6 N/P ratio) in a NanoAssemblr® Ignite instrument (Precision Nanosystems, Inc., Vancouver, Canada) at a flow rate of 8 mL/min. The LNP solution was then diluted by a factor of 3 in 25 mM Trizma^®^ hydrochloride solution (Tris-HCl, pH 7.4), dialyzed overnight against 25 mM Tris-HCl, pH 7.4 (1000X) using Slide-A-Lyzer™ G3 Dialysis Cassettes (Thermo Fisher, Massachusetts, United States). After dialysis, LNPs were concentrated using Amicon Ultra 50 K MWCO filters (Merck Millipore, Massachusetts, United States), diluted using a concentrated Tris/sucrose buffer to achieve a final concentration of 25 mM Tris/8.8% (w/v) sucrose, pH 7.4, and filtered through sterile 0.22 µm Costar® Spin-X® Centrifuge Tube Filters (Corning, New York, United States).

#### LNP Characterization

LNPs were characterized by determining hydrodynamic size (Z-average) and polydispersity index (PDI) using a Zetasizer Nano ZS (Malvern Instruments Ltd., Malvern, United Kingdom). In addition, encapsulation efficiency (EE%) and total RNA concentration were measured using a modified protocol of the Quant-iT™ RiboGreen™ RNA Assay Kit (Invitrogen, Massachusetts, United States). The Z-average size was measured to be approximately 70-90 nm and the EE% was measured to be approximately 90% for all LNPs (**Table S3**).

#### Animals

Male C57BL/6 mice, aged 8-10 weeks and sourced from Jackson Laboratory were used. Nav1.8-Cre mice (Nav1.8^Cre+^, RRID:IMSR_JAX:036564, Jackson Laboratory) or their control (Nav1.8^Cre-^) were used for infection with AAV to selectively silence the *Aak1* or *Dnm1* gene in nociceptors. Cre-dependent channel rhodopsin mice, Ai32(RCL-ChR2(H134R)/EYFP) (RRID:IMSR JAX: 012569), were crossed with the Nav1.8^Cre^ line and equal numbers of male and female offspring (Nav1.8^ChR2^) were injected intrathecally with AAV *Aak1*, *Dnm1* or *Scr* shRNA to enable optical stimulation of nociceptors. Ai14 mice (B6;129S6-*Gt(ROSA)26Sor^tm14(CAG-tdTomato)Hze^*/J) (RRID:IMSR_JAX:007914) were used to verify gene editing. Mice were housed five per cage in a temperature- (22 ± 0.5°C) and humidity-controlled *vivarium* under a controlled 12-hour light/dark cycle, with free access to food and water. At least 1 h before behavioral experiments, mice were acclimatized to the experimental room, and behavior was evaluated between 9:00 am and 5:00 pm.

#### Intrathecal injections

Mice received intrathecal (i.t.) injections between vertebrae L4 and L5 using a Hamilton syringe with 30G needle. A total volume of 5 μL was administered per mouse. For AAV delivery, Nav1.8^Cre^ or Nav1.8^ChR2^ mice were injected with 1 × 10^12^ vg/mouse. For LNP CRISPR dCas9-R experiments, C57BL/6 mice received either 300 ng or 1 µg/ 5ul of LNP formulation, and for LNP-Cre mRNA delivery, 5 µg/ 5ul was administered. A rapid tail flick was considered indicative of appropriate needle placement. Following injection, all mice presented regular motor activity consistent with that observed prior to i.t. injection.

#### Gene editing in Ai14 mice

LNP carrying the Cre recombinase mRNA were prepared as described above and were injected intrathecally into Ai14 mice. After 5 days, L4/L5 DRGs, lumbar spinal cord and brain were dissected. To identify neurons, satellite glial cells or macrophages, slides were incubated overnight at 4 °C with guinea-pig anti-NeuN (1:500, cat#AbN90, Millipore), rabbit anti-glutamine synthetase (GS, 1:5000, cat#ab49873, Abcam), or goat anti-IBA-1 (1:1000, cat#NB100-1028, Novus), respectively. Slides were washed and incubated with goat anti-rabbit Alexa Fluor 488 (1:1000; cat#A21206, Invitrogen), anti-guinea pig Alexa Fluor 488 (1:1000, cat#A11073) or anti-goat Alexa Fluor 488 (1:1000, cat#A11055) (1 hour, RT). Slides were then washed and incubated with DAPI (1 mg/mL, 5 minutes), followed by mounting in ProLong Gold Antifade (Thermo Fisher). Sections were observed using a Leica SP8 confocal microscope with HCX PL APO 20x objective or Leica DMi8 microscope equipped with a HC PL FLUOTAE 10x (NA 0.30) air objective (Leica Microsystem) and a Leica DFC9000GTC camera.

#### Collection of mouse tissue, RNAScope *in situ* hybridization and RNAScope quantification

The detailed methods were described in the supplementary material.

#### Proteome profiler array and multiplex analysis of cytokines

DRG, spinal cord and serum samples were collected at 6 h and 3 days after i.t. injection of LNP dCas9-R NC (1µg/ 5µl) or PBS in mice. The detailed methods were described in the supplementary material.

### Behavioral tests

#### Mechanical allodynia

Mechanical allodynia was assessed by measuring hindpaw withdrawal response to von Frey filament stimulation using the up-and-down method (*39*). Mice were acclimatized to the testing apparatus, which comprised individual clear Plexiglass boxes on an elevated wire mesh platform to facilitate access to the plantar surface of the hindpaws, for 1 h/d for 2 days. A series of von Frey filaments (0.02, 0.07, 0.16, 0.4, 1.0, and 2 g; Stoelting) were applied perpendicular to the plantar surface of hindpaw. The test began with an application of 0.4 g filament. A positive response was defined as a clear paw withdrawal or shaking. Whenever a positive response occurred, the next lower filament was applied, and whenever a negative response occurred, the next higher filament was applied. The testing consisted of 6 stimuli, and the pattern of response was converted to a 50% von Frey threshold(*40*). The threshold 50%, expressed in grams (g), was evaluated before (baseline) and at different time points after the treatment.

#### Heat hyperalgesia

The Hargreaves apparatus was used to evaluate hypersensitivity to heat (Ugo Basile)(*41*). Mice were acclimatized to the testing apparatus, which comprised individual clear Plexiglass chambers and a radiant heat source, for 1 h/d for 2 days. The infrared intensity was set at 50% and cut off time to a maximum of 30 seconds. The time between stimulus onset and paw withdrawal was measured automatically, giving an index of the thermal nociceptive threshold. Significant decreases in paw withdrawal latency were interpreted as evidence of thermal hyperalgesia. The latency, expressed in seconds, was evaluated before (baseline) and at different time points after the treatment.

#### Cold allodynia

Cold allodynia was assessed by measuring the acute nociceptive response to the acetone evoked evaporative cooling (*42*). A droplet (50 μL) of acetone, formed on the flat-tip needle of a syringe, was gently touched to the plantar surface of the mouse hind paw. The time spent licking and lifting of the paw over a period of 60 s was evaluated before (baseline) and at different time points after the treatment.

#### Cold hyperalgesia

Cold hyperalgesia was assessed using the cold plate apparatus(*43*). Animals were placed on a circular cold metal surface (5 °C, Hot/Cold Plate 35100, Ugo Basile Inc., Italy) enclosed by a Perspex cylinder and closely monitored to record the latency of the first nociceptive response (paw lifting, shaking, licking, or jumping). A 30 s cut-off was used to prevent potential injury to the paws. Animals were exposed only once per test day in a single trial to the cold plate. The latency, expressed in seconds, was evaluated before (baseline) and at different time points after the treatment.

#### Weight distribution

To assess the functional impact of the injury, the static weight bearing (WB) distribution was assessed in mice as previously described(*44*). Hindlimb WB was measured using a Bioseb Weight Bearing Touch; Incapacitance Test (Bioseb, Pinellas Park, FL), which measures the weight distribution across the 2 hindlimbs of a stationary animal. Mice were habituated to the testing paradigm during short sessions for 2 days prior to obtaining the pretreatment baseline measures. Three readings obtained during 10 seconds each, were collected and averaged for each animal, and the weight borne by the ipsilateral limb was expressed as a percentage of the weight borne across both hindlimbs: WB-% = (weight borne on the injured leg/weight borne on both legs) × 100%.

#### Non-evoked behavior

Non-evoked behavior was assessed using a behavioral spectrometer, which eliminates operator bias (Behavior Sequencer, Behavioral Instruments)(*10, 45*). The spectrometer comprised a 40 cm^2^ arena with a CCD camera mounted in the center of the ceiling and a door aperture in the front area of the arena. Mouse movement was assessed by a floor-mounted vibration sensor and 32 wall mounted infrared transmitter and receiver pairs. Mice were individually placed in the center of the behavioral spectrometer and their behavior was recorded, tracked, evaluated and analyzed using a computerized video tracking system (Viewer3, BiObserve) for 20 min. Average velocity of locomotion, total distance traveled (track length) in the open field, ambulation, wall distance, visits to center, grooming, locomotory activity and still time were recorded and analyzed. Non-evoked behavior was assessed 2 weeks after i.t injection of AAV *Aak1, Dnm1* or their respective *Scr* controls shRNA into Nav1.8^Cre+^ or control (Nav1.8^Cre-^) mice; or 2, 7 or 21 days after LNP dCas9-R AAK1, LNP dCas9-R Dnm1, LNP dCas9-R NC (300 ng or 1 µg/ 5µl, i.t.) or PBS (control) injection into naïve mice.

#### Locomotor activity

The motor coordination of mice was evaluated by using the Rotarod test. Mice were trained for two consecutive days before testing. The latency to fall (s) was measured with an accelerated rotation speed from 4 to 20 rpm over 3 min, 2 weeks after i.t injection of AAV *Aak1, Dnm1* or their respective *Scr* shRNA into Nav1.8^Cre+^ or control (Nav1.8^Cre-^) mice; or at 2, 3, 7, 14 and 21days after i.t. injection of LNP dCas9-R AAK1, LNP dCas9-R Dnm1, LNP dCas9-R NC (300 ng or 1 µg/ 5µl, i.t.) or PBS into naïve mice.

### Pain Models

#### Incisional pain

Postoperative pain was induced by plantar incision, as previously described(*46*). Mice were anesthetized with 2% isoflurane, and 100% O_2_ 1 L/min *via* a nose cone. After antiseptic preparation of the right hind paw with 10% povidone–iodine solution (Betadine Solution), a 5-mm longitudinal incision was made with a no. 11 blade through the skin and fascia of the plantar foot. The incision started 2 mm from the proximal edge of the heel and extended toward the toes. The skin was closed with a single mattress suture. Control mice underwent a sham procedure involving anesthesia and antiseptic preparation without an incision. AAV *Aak1*, *Dnm1* or their respective *Scr* shRNA was i.t. injected 2 weeks before plantar incision. Mechanical allodynia and heat hyperalgesia were assessed 2 hours to 7 days after the incision. LNP dCas9-R AAK1, LNP dCas9-R Dnm1, LNP dCas9-R NC (300 ng/ 5µl, i.t.) or PBS was i.t. injected immediately after the plantar incision. Mechanical allodynia and heat hyperalgesia were assessed 1 to 7 days after the incision.

#### Neuropathic pain

Spared Nerve Injury (SNI) and sham surgeries were made as described (*47*). Briefly, mice were anesthetized with isoflurane. A skin and muscle incision were made in the thigh to expose the sciatic nerve innervating the left hindpaw. The tibial and common peroneal nerves were ligated and transected distal to the ligature. The third branch, the sural nerve, was left intact. For sham controls, the nerves were exposed but not ligated or transected. AAV *Aak1*, *Dnm1* or their respective *Scr* shRNA was i.t. injected 1 week before SNI surgery. Mechanical and cold allodynia were first assessed 10 days after SNI surgery and monitored over a period of 38 days. LNP dCas9-R AAK1, LNP dCas9-R Dnm1, LNP dCas9-R NC (1µg/ 5µl, i.t.) or PBS was injected i.t. at day 10 after surgery. Mechanical and cold allodynia were assessed 1 to 28 days after the treatments.

#### Inflammatory pain

Complete Freund’s Adjuvant (CFA) (1Lmg/ml) or vehicle (0.9% NaCl) was administered by intraplantar injection (10Lµl) into the right hindpaw of sedated mice (2% isoflurane)(*10*). LNP dCas9-R AAK1, LNP dCas9-R Dnm1, LNP dCas9-R NC (300 ng/ 5µl, i.t.) or PBS was injected i.t. 24 h after CFA. Mechanical allodynia and heat hyperalgesia were assessed 1 to 13 days after the treatments.

#### Osteoarthritis pain

Mice were anesthetized (2% isoflurane) and received intra-articular injection of monosodium iodoacetate (MIA, 1 mg/mL; cat# 57858-25G-F; Milipore Sigma) through the infra-patellar ligament of the left knee(*48*). Mechanical allodynia and weight distribution were assessed 10 days after MIA injection and monitored for 28 days. LNP dCas9-R AAK1, LNP dCas9-R Dnm1, LNP dCas9-R NC (1 µg/5 µl, i.t.), or PBS was injected i.t. at day 10 after MIA. Mechanical allodynia and weight distribution were assessed 1 to 28 days after the treatments.

### Electrophysiology

#### Spinal slice preparation

At least 2 weeks following AAV i.t. injection, adult mice were anesthetized (5% isoflurane), decapitated and the lumbar region of the spinal cord with the dorsal root exposed by laminectomy was removed. Parasagittal spinal cord slices with the dorsal root attached (300 μm) were sectioned on a vibratome (Leica VT 1200s) in ice cold (0-4°C) oxygenated sucroseLbased ACSF that contained (mM): 100 sucrose, 63 NaCl, 2.5 KCl, 1.2 NaH2PO4, 1.2 MgCl2, 25 glucose, 25 NaHCO3 and 5 Na ascorbate. Slices were then incubated for 15 min at 34°C in NMDGLbased recovery ACSF composed of (mM): 93 NMDG, 2.5 KCl, 1.2 NaH2PO4, 30 NaHCO3, 20 HEPES, 25 glucose, 5 Na ascorbate, 2 thiourea, 3 Na pyruvate, 10 MgSO4 and 0.5 CaCl2 and adjusted to pH 7.4 with HCl. After the recovery incubation, slices were transferred to oxygenated ACSF with the following composition (mM): 125 NaCl, 2.5 KCl, 1.25 NaH2PO4, 1.2 MgCl2, 2.5 CaCl2, 25 glucose and 25 NaHCO3 for 45 min at 36°C and then maintained at RT prior to transfer to the recording chamber. All ACSF solutions were equilibrated with 95% O2 and 5% CO2.

#### Spinal cord electrophysiology

Slices were transferred to the recording chamber and continuously superfused with ACSF equilibrated with 95% O2/5% CO2 at a rate of 2ml/min at RT. Dodt-contrast optics were used to identify dorsal horn neurons in the translucent substantia gelatinosa layer of the superficial dorsal horn. Optogenetically-evoked post-synaptic currents (oPSCs) were recorded in whole-cell voltage clamp using a CsCl-based internal solution composed of (mM): 140 CsCl, 10 EGTA, 5 HEPES, 2 CaCl2, 2 MgATP, 0.3 NaGTP, 5 QX-314.Cl and 0.1% biocytin (osmolarity 285–295 mosmol/l). Patch clamp electrodes had resistances between 5-10 MΩ and neurons were held at -65 mV (not corrected for the liquid junction potential of 4 mV). An optical probe tip (100 um, NA 0.66, with a 45° angle) was placed in the dorsal root entry zone for optical stimulation using a 465 nm LED light source (Doric lenses, Quebec). Electrophysiological data was recorded in pClamp 11 (Molecular Devices). Optically evoked excitatory synaptic currents (oEPSCs) were recorded in the presence of gabazine (10 μM) and strychnine (0.5 μM) and to measure the contribution of SP and CGRP, oEPSCs were recorded in the presence of Aprepitant (1 μM) and Olcegepant (1 μM). Glutamate-mediated currents were blocked with NBQX (10 μM), AP5 (100 μM) at the end of the experiment to confirm that remaining currents were AMPA and NMDA-mediated.

#### Statistical Analysis

The results are expressed as the mean ± SEM. Differences were assessed using Student 2-tailed *t* test for 2 comparisons and 1- or 2-way ANOVA with Sídák, Tukey, Newman-Keuls, or Dunnett post hoc test for multiple comparisons. Statistical analyses were performed on raw data using GraphPad Prism 8 (GraphPad Software Inc.). P-values less than 0.05 (P < 0.05) were considered significant. The statistical tests used and sample size for each analysis are shown in the Figure legends.

## List of Supplementary Materials

Fig. S1 to S7

Table S1 to S3

## Funding

National Institutes of Health grant R01NS102722 (NWB)

National Institutes of Health grant R01DE029951 (NWB)

Department of Defense grant W81XWH2210238 (NWB)

Australian Research Council grant ARC DP190102854 (WLI)

Finanziata con il contributo del Ministero dell’Università e della ricerca ai sensi del D.D. n.1236. del 1° agosto 2023 – BANDO FIS 2 (FDL, RN)

## Author contributions

Conceptualization: RT, NWB, CWP, WLI

Methodology: RT, NWB, CWP, WLI, FDL, RN

Investigation: RT, MFPF, TP, ED, MC

Visualization: RT, NWB, MFPF

Funding acquisition: NWB, CWP, WLI

Project administration: RT, NWB

Supervision: RT, NWB

Writing - original draft: RT, NWB

Writing - review & editing: NWB, CWP, WLI, FDL, RN

## Competing interests

N. W. Bunnett is a founding scientist of Endosome Therapeutics Inc. Research in NW Bunnett laboratory is partly supported by Takeda Pharmaceuticals Inc. The remaining authors have no conflicts of interest to declare.

## Data and materials availability

Contact the corresponding author (R. Tonello at) to obtain original data.

## Notes

### Competing Interest Statement

NW Bunnett is a founding scientist of Endosome Therapeutics Inc. Research in NW Bunnett laboratory is partly supported by Takeda Pharmaceuticals Inc. The remaining authors have no conflicts of interest to declare.

