## Supplemental Material for "Targeting Synaptic Vesicle Endocytosis in Nociceptors Provides Sustained Pain Relief"

**Supplementary Methods**

**Collection of mouse tissue.** Mice were anesthetized (5% isoflurane) and perfused through the ascending aorta with PBS and then 4% paraformaldehyde in PBS. DRG (L4-L5), spinal cord and brain were removed, fixed in 4% paraformaldehyde in PBS at 4°C for 1 hour or overnight, respectively, cryoprotected in 30% sucrose (24 h, 4°C), and embedded in Optimal Cutting Temperature compound (Tissue Tek). Frozen sections (10-12 μm) were mounted onto Superfrost Plus slides (Fisher), dried (15 min) and stored (-20°C).

**RNAScope *in situ* hybridization.** The RNAScope system (Advanced Cell Diagnostics) was used per the manufacturer’s directions for fresh-frozen tissue except for the omission of the initial on-slide fixation step. Probe hybridization and detection using the Multiplex Fluorescent Kit v2 followed the manufacturer’s directions. Probes to *Mm-Aak1* (#1097711-C1), *Mm-Dnm1* (#446931-C3) and *Mn-Scn10a* (#426011-C2) were used. Sections were incubated with TSA VividTM Fluorophore 570 (1:1500, cat#323272, Advanced Cell Diagnostics) or TSA VividTM Fluorophore 650 (1:1500, cat#323273) for fluorescence detection. Slides were washed and incubated with DAPI, 4′,6-diamidino-2-phenylindole (1 µg/ml, 5 min) and mounted in ProLong® Gold Antifade. Sections were observed using a Leica SP8 confocal microscope with HCX PL APO 20x objective and 40x oil objective.

**RNAScope quantification.** *Aak1* and *Dnm1* mRNA were localized by RNAScope. Confocal images were analyzed using Fiji ImageJ (NIH) according to ACD Bio-Techne Technical Note. *AAV approach:* Regions of interest corresponding to AAK1 or Dnm1 expression were defined by applying a threshold with the Moments setting (Min & Max) and analyzing particles with sizes ranging from 0 to infinity. Regions of interest were overlaid on the *Scn10a* mRNA-positive or -negative (Nav1.8+ve or Nav1.8-ve neurons) micrograph, which were previously defined by applying a threshold with Li setting. The number of dots per Nav1.8+ve and -ve neurons was quantified. Results are expressed as dots/Nav1.8+ or Nav1.8- neurons. *LNP approach:* Regions of interest corresponding to AAK1 or Dnm1 expression were defined by applying a threshold with the Moments setting (Min & Max) and analyzing particles with sizes ranging from 0 to infinity. Regions of interest were overlaid on the original micrograph and the number of dots per area was quantified. Results are expressed as dots/µm^2^ tissue. A total of 3 images (20X magnification) were analyzed for each mouse (N=4-7 mice for control and treatment groups; at least 12 images were analyzed per experimental group).

**Proteome profiler array.** DRG and spinal cord samples were collected at 6 h and 3 days after i.t. injection of LNP dCas9-R NC (1µg/ 5µl) or PBS in mice for a cytokine array analysis using the Proteome Profiler Mouse Cytokine Array Kit, Panel A (ARY006, R&D Systems) according to manufacturer’s instructions. Tissue lysates were prepared in 1% Triton X-100 in PBS containing a protease inhibitor cocktail (Aprotinin, Leupeptin, and Pepstatin, all 10 µg/ml) at 4 °C. Signal was developed using an imaging system (ChemiDoc, Bio-Rad). The density of specific cytokine dots was measured using Fiji ImageJ (NIH).

**Multiplex analysis of cytokines.** Serum was collected at 6 h and 3 days after i.t. injection of LNP dCas9-R NC (1µg/ 5µl) or PBS in mice. Thirty-six cytokines, chemokines and growth factors were measured using the Eve Technologies’ Mouse Cytokine/Chemokine 36-Plex Discovery Assay Array (MD36) as per the manufacturer's instructions. The multiplexing analysis was performed by Eve Technologies Corporation (Calgary, Alberta, Canada) using the Luminex® 200™ system (Luminex Corporation/DiaSorin, Saluggia, Italy) with Bio-Plex Manager™ software (Bio-Rad Laboratories Inc., Hercules, California, USA).

**Supplementary Figures**

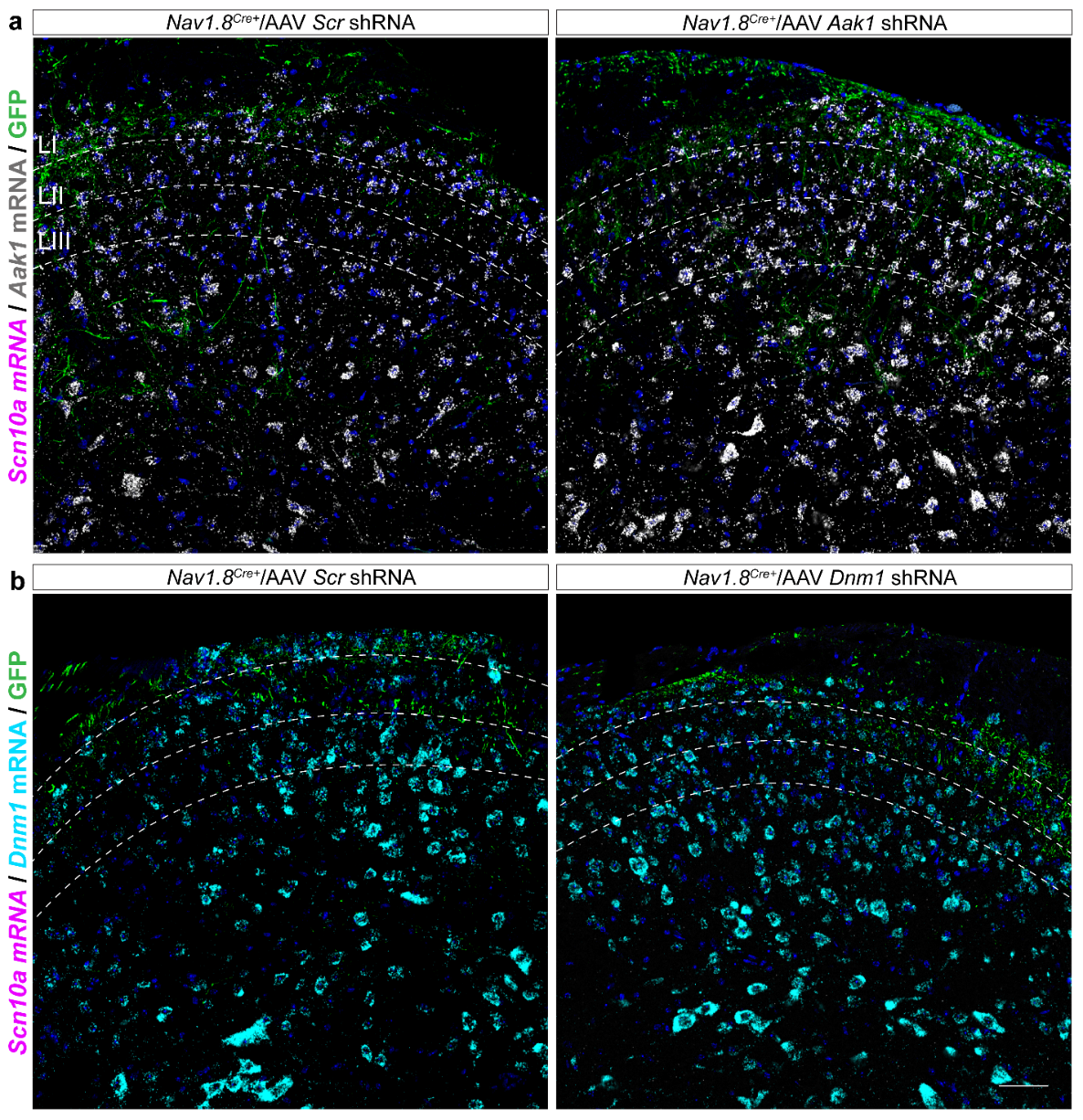

**Fig. S1. Expression of *Aak1* and *Dnm1* mRNA in spinal cord after AAV shRNA injection in Nav1.8^Cre+^ mice.** Representative images comparing the expression of *Aak1* mRNA (a) and *Dnm1* mRNA (b) in the spinal cord 3 weeks after intrathecal injection of AAV *Aak1*, *Dnm1*, or their respective *scrambled* controls (*Scr)* shRNA in *Nav1.8^Cre+^* mice. n=4 mice per group. GFP-positive projections from Nav1.8+ve neurons. Scale bar: 50 µm.

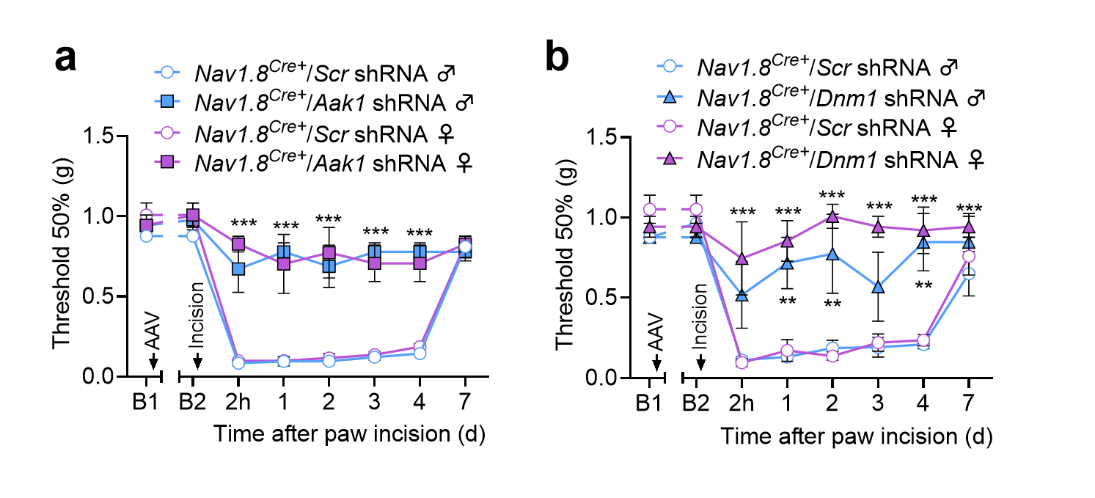

**Fig. S2. Selective knockdown of AAK1 or Dnm1 in nociceptors prevents postoperative pain in male and female mice.** Mechanical allodynia of the plantar incision in *Nav1.8^Cre^* male and female mice, measured from 2 hours to 7 days after incision. Mice received intrathecal (i.t.) injections of AAV *Aak1* shRNA (a), AAV *Dnm1* shRNA(b) or their respective *scrambled* controls (*Scr)* shRNA 2 weeks prior to the incision, n=3-4 mice per group. Data are presented as Mean ± SEM. **P<0.01, ***P<0.001 vs. *Nav1.8^Cre+^/Scr* shRNA. Two-way ANOVA with Sídák multiple comparisons test. B1: baseline 1, pre-AAV injection; B2: baseline 2, two weeks post-AAV injection.

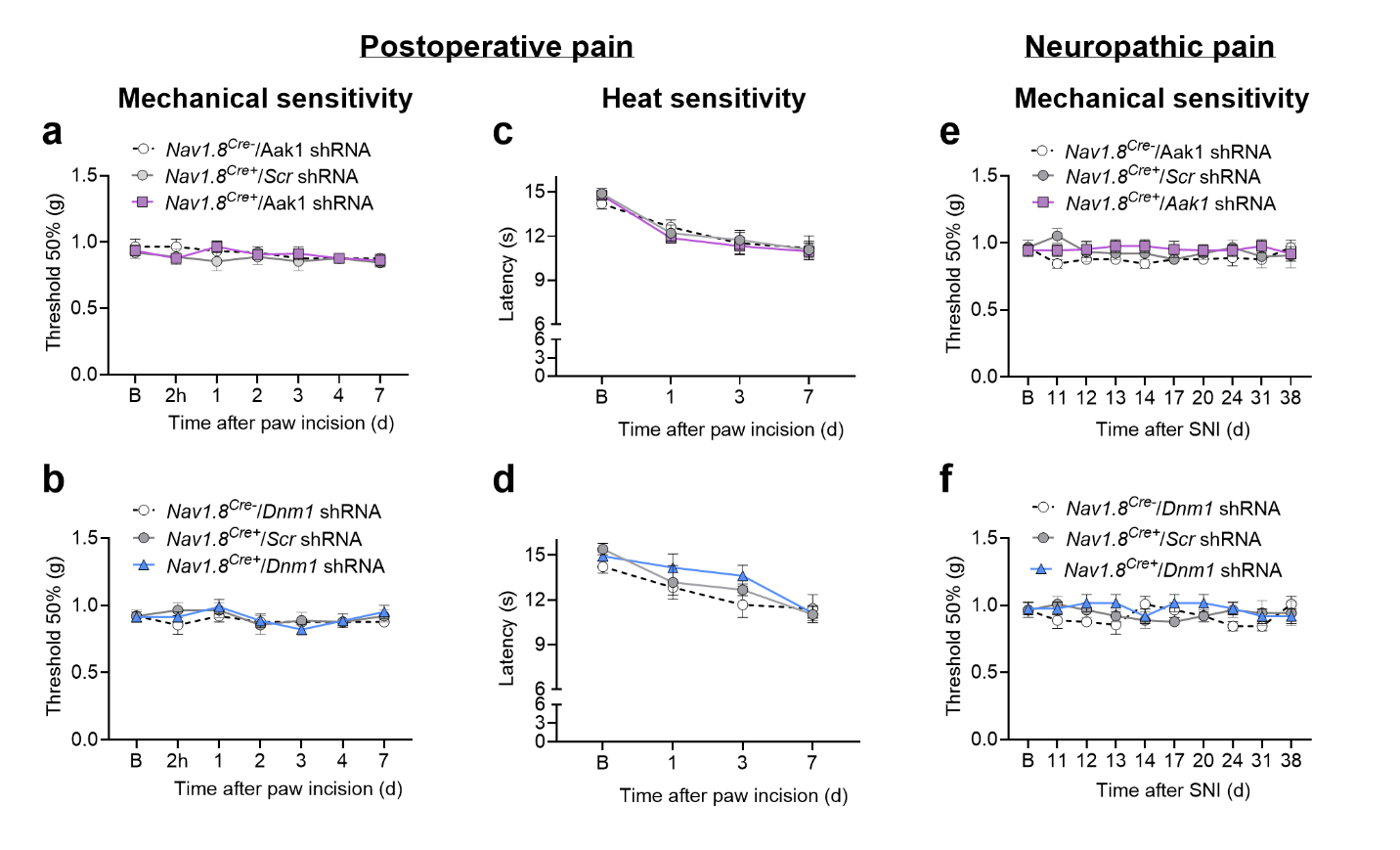

**Fig. S3. Effect of selective knockdown of AAK1 or Dnm1 in nociceptors on mechanical and thermal sensitivity in the contralateral paw on postoperative and neuropathic pain models.** Time course of the mechanical (a, b, e, f) and heat sensitivity (c, d) in the contralateral hind paw after plantar incision (a-d) or 10 days after SNI surgery (e, f) to the ipsilateral paw in *Nav1.8^Cre^* male and female mice. Mice received intrathecal injections of AAV *Aak1*, *Dnm1*, or their respective *scrambled* controls (*Scr)* shRNA two weeks prior to the incision and one week prior to the SNI surgery, n=6-8 mice per group. 2-way ANOVA, Sídák multiple comparisons test.

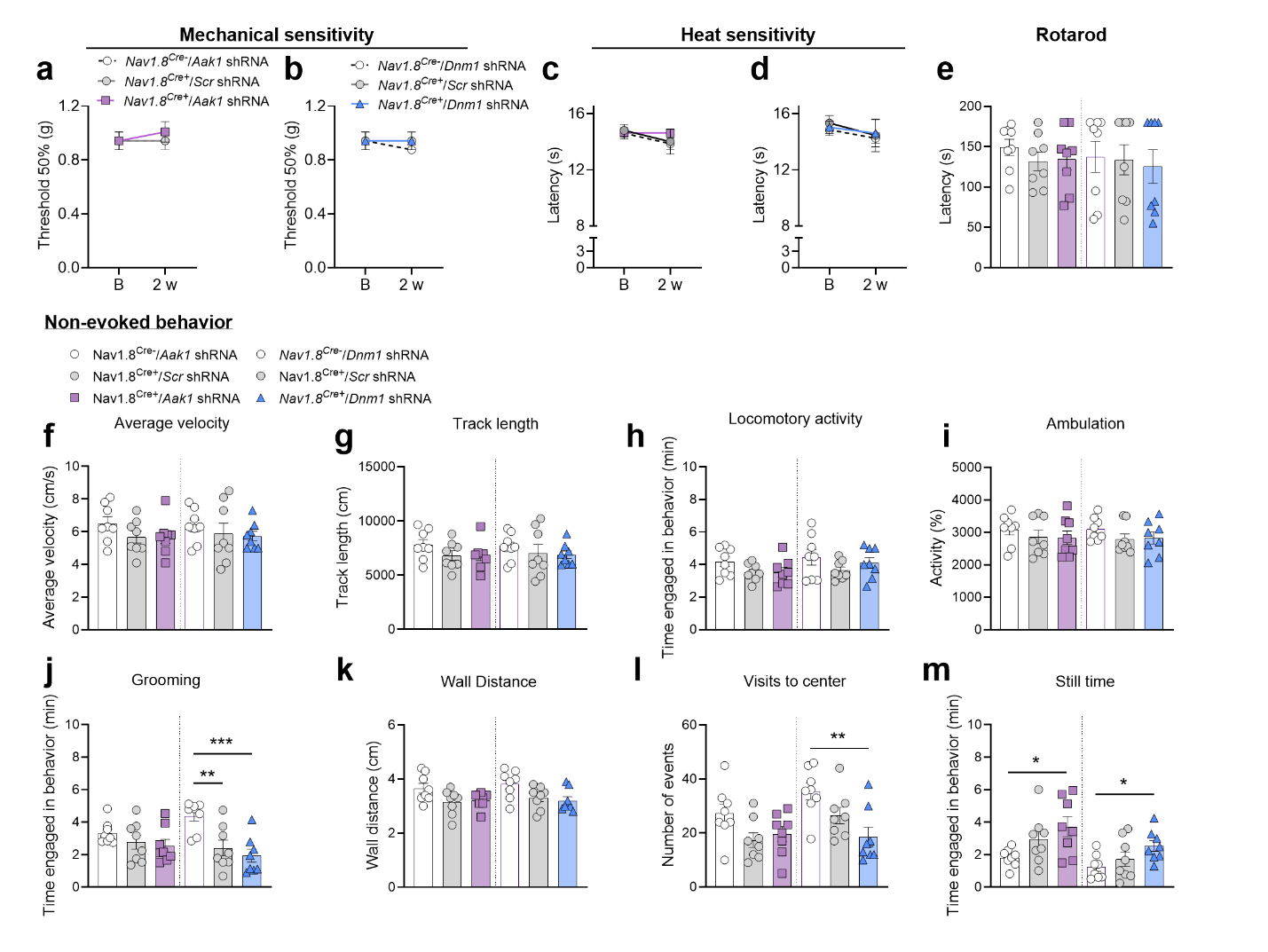
**Fig. S4. Sensitive, locomotor, exploratory and grooming behaviors.** Mechanical (a, b) and heat (c, d) sensitivity, locomotor activity (rotarod) (e) and non-evoked behavior (f-m) at 2 weeks after intrathecal injection of AAV *Aak1*, *Dnm1*, or their respective *scrambled* controls (*Scr)* shRNA in *Nav1.8^Cre^* male and female mice. n = 8 mice per group. Data are presented as Mean±SEM. *P<0.05, **P<0.01, ***P<0.001 vs. *Nav1.8^Cre-^*. 1-way ANOVA, Tukey multiple comparisons test. 2 w: 2 weeks.

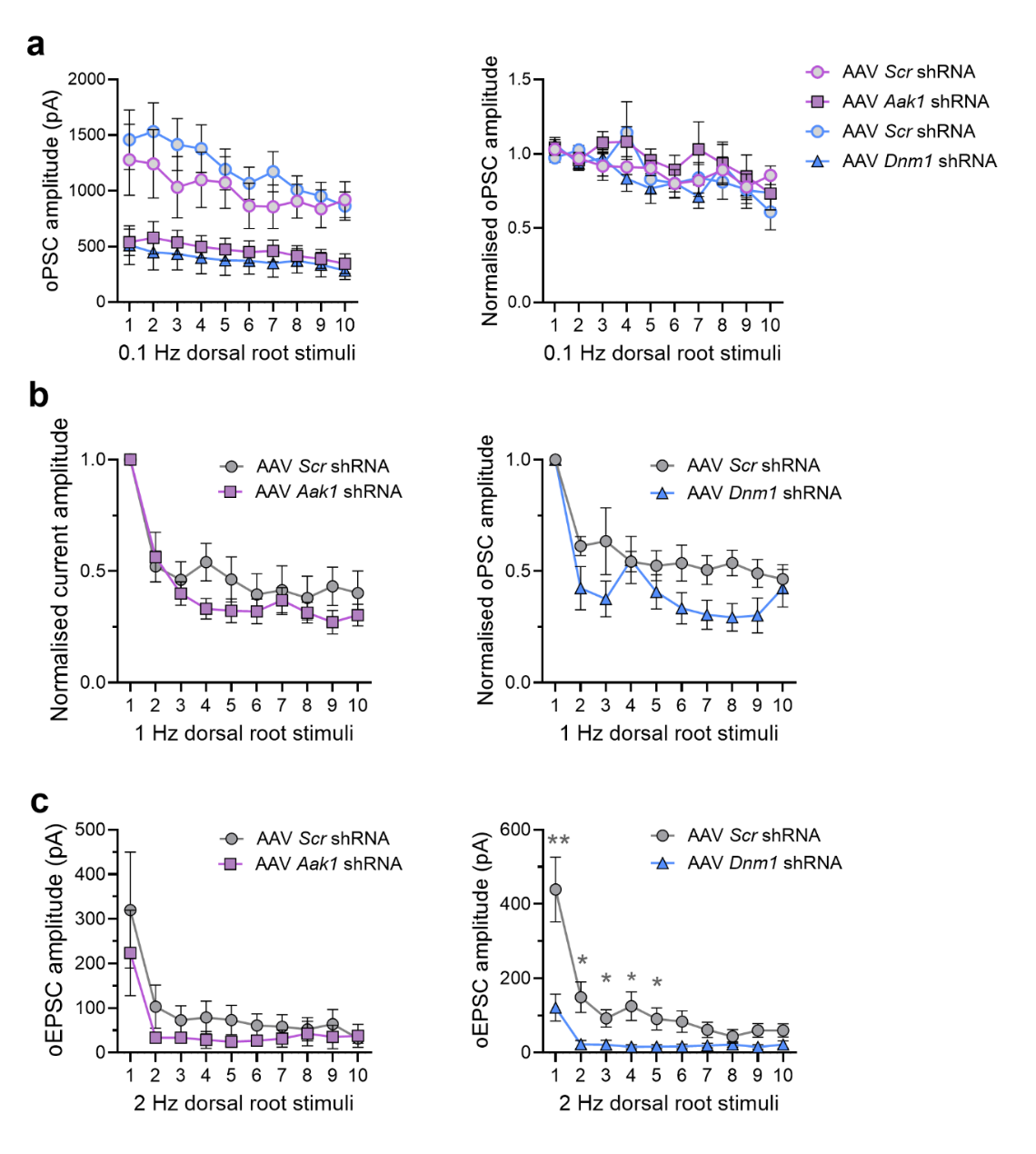

**Fig. S5. Effect of increasing stimulus frequency on synaptic current amplitude following selective knockdown of AAK1 and Dnm1 in nociceptors.** Mean and normalized oPSC amplitude in response to 0.1Hz stimuli. Current amplitudes were normalized to the mean of the first two pulses (a). Normalized oPSC amplitude in response to 1Hz stimuli (normalized to the first pulse) (b). Mean oEPSC amplitude in response to 2Hz stimuli (c). Mean±SEM. *P<0.05, **P<0.01 versus control. Unpaired t-test.

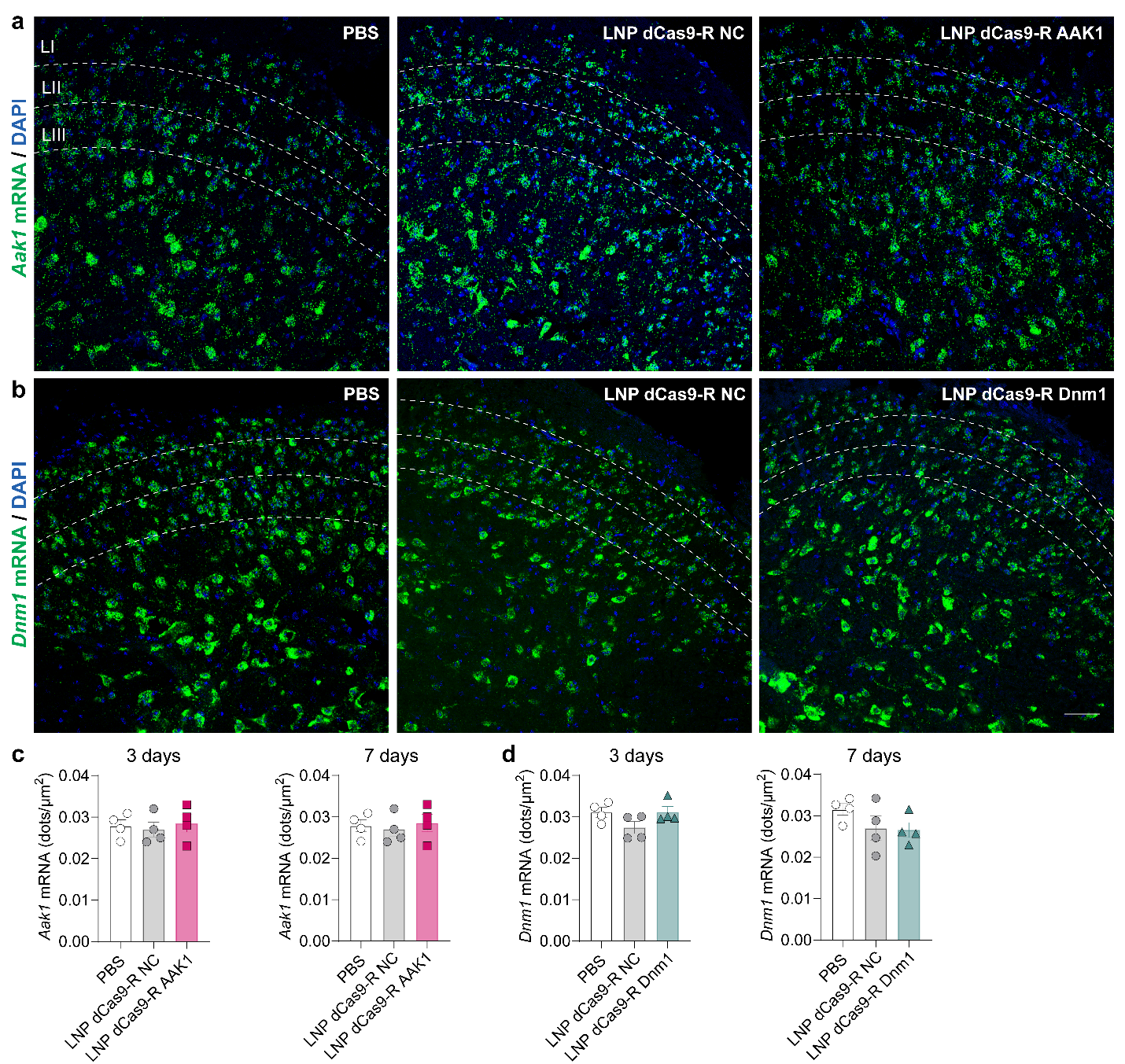

**Fig. S6. Expression of *Aak1* and *Dnm1* mRNA in spinal cord after LNP-encapsulated CRISPR/dCas9-R AAK1 or Dnm1 injection.** RNAScope localization (a, b) quantification (number of dots per area) (c, d) of *Aak1* (a, c) and *Dnm1* (b, d) *mRNA* expression in mouse spinal cord after intrathecal injection of LNP dCas9-R AAK1, LNP dCas9-R Dnm1, LNP dCas9-R NC or PBS at days 3 (300 ng/ 5µl, i.t.) and 7 (1 µg/ 5µl, i.t.) post treatment, n= 4 mice per group. Scale bar: 50 µm.

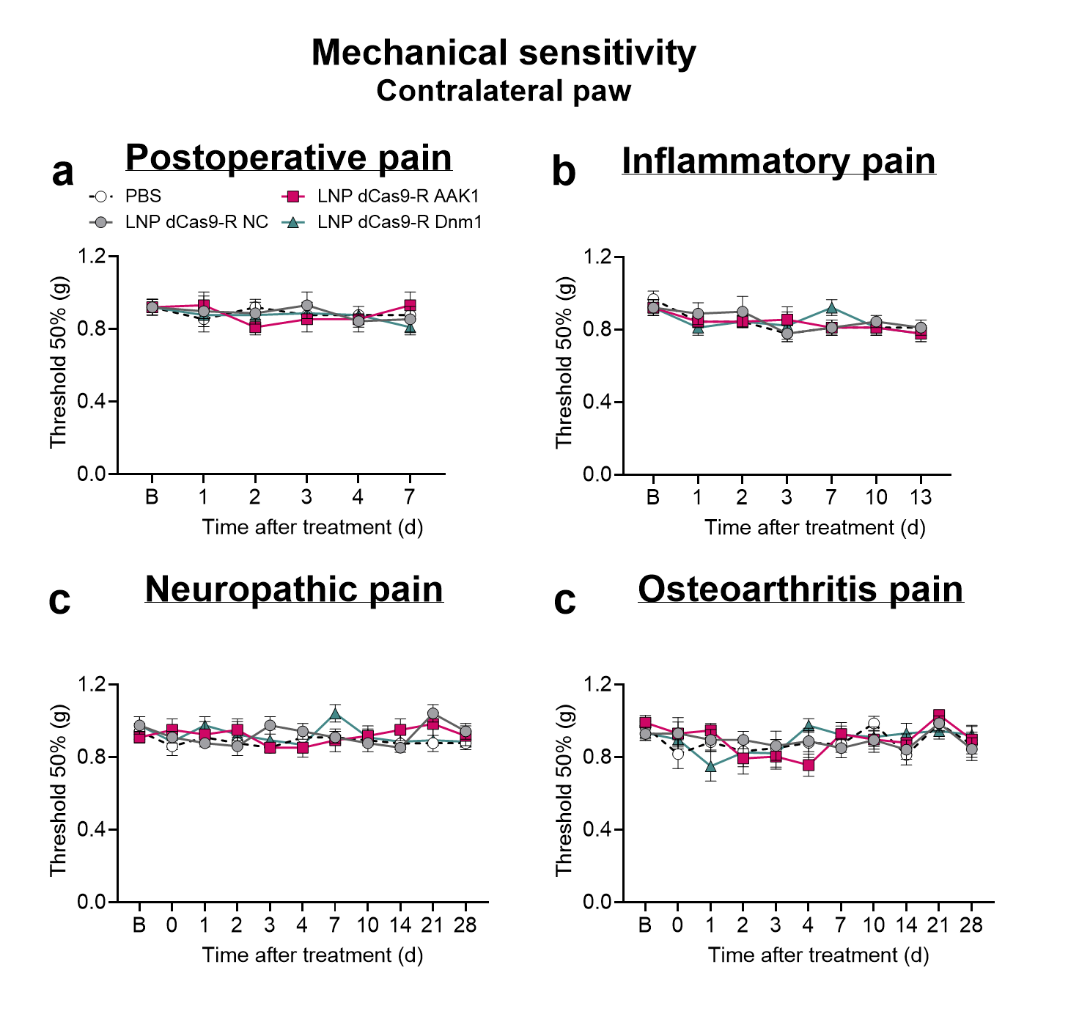

**Fig. S7. Effect of LNP-encapsulated CRISPR/dCas9-R AAK1 or Dnm1 on mechanical sensitivity in the contralateral paw on postoperative, inflammatory, neuropathic and osteoarthritis pain models.** Time course of the effect of LNP dCas9-R AAK1, LNP dCas9-R Dnm1, LNP dCas9-R NC (300 ng or 1 µg/ 5µl, i.t.) or PBS on mechanical sensitivity tested in the contralateral hind paw after plantar incision (a), Complete Freund’s Adjuvant (CFA) intraplantar injection (b), Spared Nerve Injury (SNI) surgery (c) or monosodium iodoacetate (MIA) intra-articular injection (d) to the ipsilateral paw in mice, n=6-8 mice per group. 2-way ANOVA, Sídák multiple comparisons test.

**Supplementary Tables**

**Table S1: Primers for plasmid construction**

| PRIMER | SEQUENCE 5’-3’ |
| --- | --- |
| P1 | ggcaGACCCTATTCCTGTACTAATTATACATCTGTGGCTTCACTATAATTAGTACAGGAATAGGGTT |
| P2 | agcgAACCCTATTCCTGTACTAATTATAGTGAAGCCACAGATGTATAATTAGTACAGGAATAGGGTC |
| P3 | ggcaTCCTGCGACATTCTATAAATATTACATCTGTGGCTTCACTAATATTTATAGAATGTCGCAGGG |
| P4 | agcgCCCTGCGACATTCTATAAATATTAGTGAAGCCACAGATGTAATATTTATAGAATGTCGCAGGA |
| P5 | ggcaGTCATAGCCTACTCCTTATATATACATCTGTGGCTTCACTATATATAAGGAGTAGGCTATGAT |
| P6 | agcgATCATAGCCTACTCCTTATATATAGTGAAGCCACAGATGTATATATAAGGAGTAGGCTATGAC |
| P7 | ggcaGCTTGCTTAAACTAATCCTATATACATCTGTGGCTTCACTATATAGGATTAGTTTAAGCAAGT |
| P8 | agcgACTTGCTTAAACTAATCCTATATAGTGAAGCCACAGATGTATATAGGATTAGTTTAAGCAAGC |

**Table S2: sgRNA sequences**

| **sgRNA sequence name** | **sgRNA spacer sequence** |
| --- | --- |
| MM_AAK1_Ci_sg1 | AGGGGUCCGGUGCGUUUCCA |
| MM_AAK1_Ci_sg2 | GGCCUGCGACACGAAGGAGG |
| MM_AAK1_Ci_sg3 | GCGCUCGGGGAUCAGCUGGC |
| MM_Dnm1_Ci_sg1 | GCGCACGCAGUCGGGAUAUU |
| MM_Dnm1_Ci_sg2 | GGGCCCGGCGGGCUGCGAUC |
| MM_Dnm1_Ci_sg3 | GCCGGGCCCGCCGCAACCAU |
| NC sgRNA | UCAUGCUUGCUUGGGCAAAA |

**Table S3: Lipid nanoparticle characterization data**

| **LNP name** | **Batch Number** | **Particle Size**  **(Z-average, nm)** | **Polydispersity index (PDI)** | **Encapsulation Efficiency (EE%)** |
| --- | --- | --- | --- | --- |
| dCas9-R AAK1 | 1 | 79.73 | 0.121 | 90.0 |
| dCas9-R AAK1 | 2 | 83.73 | 0.117 | 95.3 |
| dCas9-R AAK1 | 3 | 91.30 | 0.111 | 95.5 |
| dCas9-R Dnm1 | 1 | 73.84 | 0.138 | 94.1 |
| dCas9-R Dnm1 | 2 | 79.66 | 0.144 | 94.7 |
| dCas9-R Dnm1 | 3 | 77.70 | 0.070 | 96.6 |
| dCas9 NC | 1 | 77.64 | 0.100 | 88.9 |
| dCas9 NC | 2 | 75.50 | 0.070 | 96.7 |
| Cre recombinase | 1 | 83.30 | 0.112 | 94.2 |
| Cre Recombinase | 2 | 77.90 | 0.090 | 96.4 |
